# An open, analysis-ready, and quality controlled resource for pediatric brain white-matter research

**DOI:** 10.1101/2022.02.24.481303

**Authors:** Adam Richie-Halford, Matthew Cieslak, Lei Ai, Sendy Caffarra, Sydney Covitz, Alexandre R. Franco, Iliana I. Karipidis, John Kruper, Michael Milham, Bárbara Avelar-Pereira, Ethan Roy, Valerie J. Sydnor, Jason Yeatman, The Fibr Community Science Consortium, Theodore D. Satterthwaite, Ariel Rokem

## Abstract

We created resources to facilitate research on the role of human brain microstructure in the development of mental health disorders, based on openly-available diffusion MRI (dMRI) data from the Healthy Brain Network (HBN) study. First, we curated the HBN dMRI data (N=2747) into the Brain Imaging Data Structure and preprocessed it according to best-practices, including denoising and correcting for motion effects, susceptibility-related distortions, and eddy currents. Preprocessed, analysis-ready data was made openly available. Data quality plays a key role in the analysis of dMRI, and we provide automated quality control (QC) scores for every scan, as part of the data release. To scale QC to this large dataset, we trained a neural network through the combination of a small data subset scored by experts and a larger set scored by community scientists. The network performs QC highly concordant with that of experts on a held out set (ROC-AUC = 0.947). A further analysis of the neural network demonstrates that it relies on image features with relevance to QC. Altogether, this work both delivers a resource for transdiagnostic research in brain connectivity and pediatric mental health and serves as a novel tool for automated QC of large datasets.

## Introduction

Childhood and adolescence are characterized by rapid dynamic change to human brain structure and function (***Lebel and Deoni, 2018***). This period of development is also a time during which the symptoms of many mental health disorders emerge (***Paus et al., 2008***). Understanding how individual differences in brain development relate to the onset and progression of psychopathology inevitably requires large datasets (***Paus, 2010***; ***Fair et al., 2021***). The Healthy Brain Network (HBN) is a landmark pediatric mental health study that is designed to eventually include MRI images along with detailed clinical and cognitive phentoyping from over 5000 New York City area children and adolescents (***Alexander et al., 2017***). The HBN dataset takes a trans-diagnostic approach and provides a broad range of phenotypic and brain imaging data for each individual. One of the brain imaging measurements acquired is diffusion MRI (dMRI), which is the dominant technology for inferring the physical properties of white matter (***Wandell, 2016***). The dMRI data is openly available in its raw form through the Functional Connectomes Project and the International Neuroimaging Data-Sharing Initiative (FCP-INDI), spurring collaboration on open and reproducible science (***Mennes et al., 2013***). However, this raw, publicly available data requires extensive processing and quality assurance before it can be fruitfully analyzed.

The analysis of a large, multi-site dMRI dataset must take into account the inevitable variability in scanning parameters across scanning sessions. Critical preprocessing steps, such as susceptibility distortion correction (***Jones and Cercignani, 2010***) require additional MRI acquisitions besides dMRI and accurate metadata accompanying each image. A session missing an acquisition or important metadata can either be processed to the extent its available data allows or excluded entirely. In addition, the quality of preprocessed data is heavily affected by differences in acquisition parameters (***Yeh et al., 2019***) and by differences in preprocessing steps. Here we address these problems by meticulously curating the HBN data according to the Brain Imaging Data Specification (BIDS) (***Gorgolewski et al., 2016***) and processing the data using the *QSIPrep* (***Cieslak et al., 2021***) BIDS App (***Gorgolewski et al., 2017***). *QSIPrep* automatically builds and executes benchmarked workflows that adhere to best practices in the field given the available BIDS data. The results include automated data quality metrics, visual reports and a description of the processing steps automatically chosen to process each session.

This preprocessing requires a costly compute infrastructure and is both time-consuming and error-prone. Requiring researchers to process dMRI data on their own introduces both a practical barrier to access and an extra source of heterogeneity into the data, devaluing its scientific utility. We provide the preprocessed data as a transparent and open resource, thereby reducing barriers to data access and allowing researchers to spend more of their time answering questions in brain development and psychopathology rather than recapitulating preprocessing.

In addition to requiring extensive preprocessing, dMRI data must be thoroughly checked for quality. dMRI measurements are susceptible to a variety of artifacts that affect the quality of the signals and the ability to make accurate inferences from them. In small studies, with few participants, it is common to thoroughly examine the data from every participant as part of a quality control (QC) process. However, expert examination is time consuming and is prohibitive in large datasets such as HBN. This difficulty could be ameliorated through the automation of QC. Given their success in other visual recognition tasks, machine learning and computer vision methods, such as convolutional deep artificial neural networks or “deep learning” (***LeCun et al., 2015***), are promising avenues for automation of QC. However, one of the challenges of these new methods is that they require a large training dataset to attain accurate performance. In previous work, we demonstrated that deep learning can accurately emulate expert QC of T1-weighted (T1w) anatomical brain images (***Keshavan et al., 2019***). To obtain a large enough training dataset of T1w images in our prior study, we deployed a community science tool ^1^ that collected quality control scores of parts of the dataset from volunteers through a web application. The scores were then calibrated using a gold standard expert-scored subset of these images. A deep learning neural network was trained on the calibrated and aggregated score, resulting in very high concordance with expert ratings on a separate test dataset. We termed this approach “hybrid QC”, because it combined information from experts with information from community scientists to create a scalable machine learning algorithm that can be applied to future data collection.

However, the hybrid QC proof-of-concept left lingering questions about its applicability to other datasets because it was trained on a single-site, single-modality dataset. Here, we expand the hybrid-QC approach to a large multi-site dMRI dataset. Moreover, one of the common critiques of deep learning is that it can learn irrelevant features of the data and does not provide information that is transparent enough to interpret (***Lipton, 2017***; ***Salahuddin et al., 2022***; ***Zech et al., 2018***). To confirm that the hybrid-QC deep learning algorithm uses meaningful features of the diffusion-weighted images (DWI) to perform accurate QC, we used machine learning interpretation methods that pry open the “black box” of the neural network, thereby highlighting the features that lead to a specific QC score (***Sundararajan et al., 2017***; ***Murdoch et al., 2019***).

Taken together, the combination of curated BIDS data, preprocessed images, and quality control scores generated by the deep learning algorithm provides researchers with a rich and accessible data resource. We anticipate that these HBN Preprocessed Open Diffusion Derivatives (HBN-POD2) will accelerate translational research on both normal and abnormal brain development.

## Results

The aims of this study were fourfold: (i) curate the HBN MRI data into a fully BIDS-compliant MRI dataset, (ii) perform state-of-the-art diffusion MRI (dMRI) preprocessing using *QSIPrep*, (iii) assign QC scores to each participant, and (iv) provide unrestricted public release to the outputs from each of these steps. We started with MRI data from 2747 HBN participants available through FCP-INDI, curating these data for compliance with the Brain Imaging Data Structure (BIDS) Specification (***Gorgolewski et al., 2016***). We preprocessed the structural MRI (sMRI) and diffusion MRI (dMRI) data using *QSIPrep*. Participants that could not be curated to comply with the BIDS standard or that did not have dMRI data were excluded, resulting in 2134 participants with preprocessed, BIDS-compliant dMRI data (Figure 1). HBN neuroimaging data was collected at four sites: Staten Island (SI, *N* = 300), Rutgers University Brain Imaging Center (RU, *N* = 873), the CitiGroup Cornell Brain Imaging Center (CBIC, *N* = 887), and the City University of New York Advanced Science Research Center (CUNY, *N* = 74), where numbers in parentheses represent participant counts in HBN-POD2. Figure 2 depicts the age distribution of study participants by sex for each of these scan site as well as pairwise distributions for the automated quality metrics that are described in the next section.

**Figure 1.**
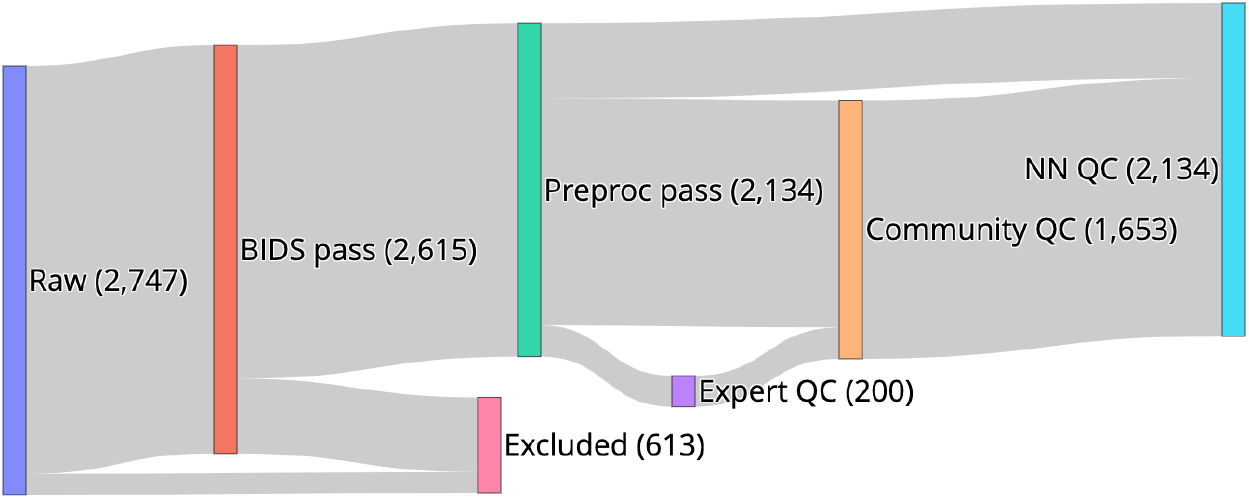
HBN-POD2 data provenance: Imaging data for 2747 participants, aged 5-21 years and collected at four sites in the New York City area, was made available through the Functional Connectomes Project and the International Neuroimaging Data-Sharing Initiative (FCP-INDI). These data were curated for compliance to the BIDS Specification (***Gorgolewski et al., 2016***) and availability of imaging metadata in json format. 2615 participants met this Specification. Imaging data was preprocessed using *QSIPrep* (***Cieslak et al., 2021***) to group, distortion correct, motion correct, denoise, coregister and resample MRI scans. Of the BIDS curated participants, 2134 passed this step, with the majority of failures coming from participants with missing dMRI scans. Expert raters assigned QC scores to 200 of these participants, creating a “gold standard” QC subset. Community raters then assigned binary QC ratings to a superset of the gold standard containing 1653 participants. An image classification algorithm was trained on a combination of automated quality metrics from *QSIPrep* and community scientist reviews to “extend” the expert ratings to the community science subset. Finally, a deep learning QC model was trained on the community science subset to assign QC scores to the entire dataset and to future releases from HBN. The HBN-POD2 dataset, including QC ratings, is openly available through FCP-INDI.

**Figure 2.**
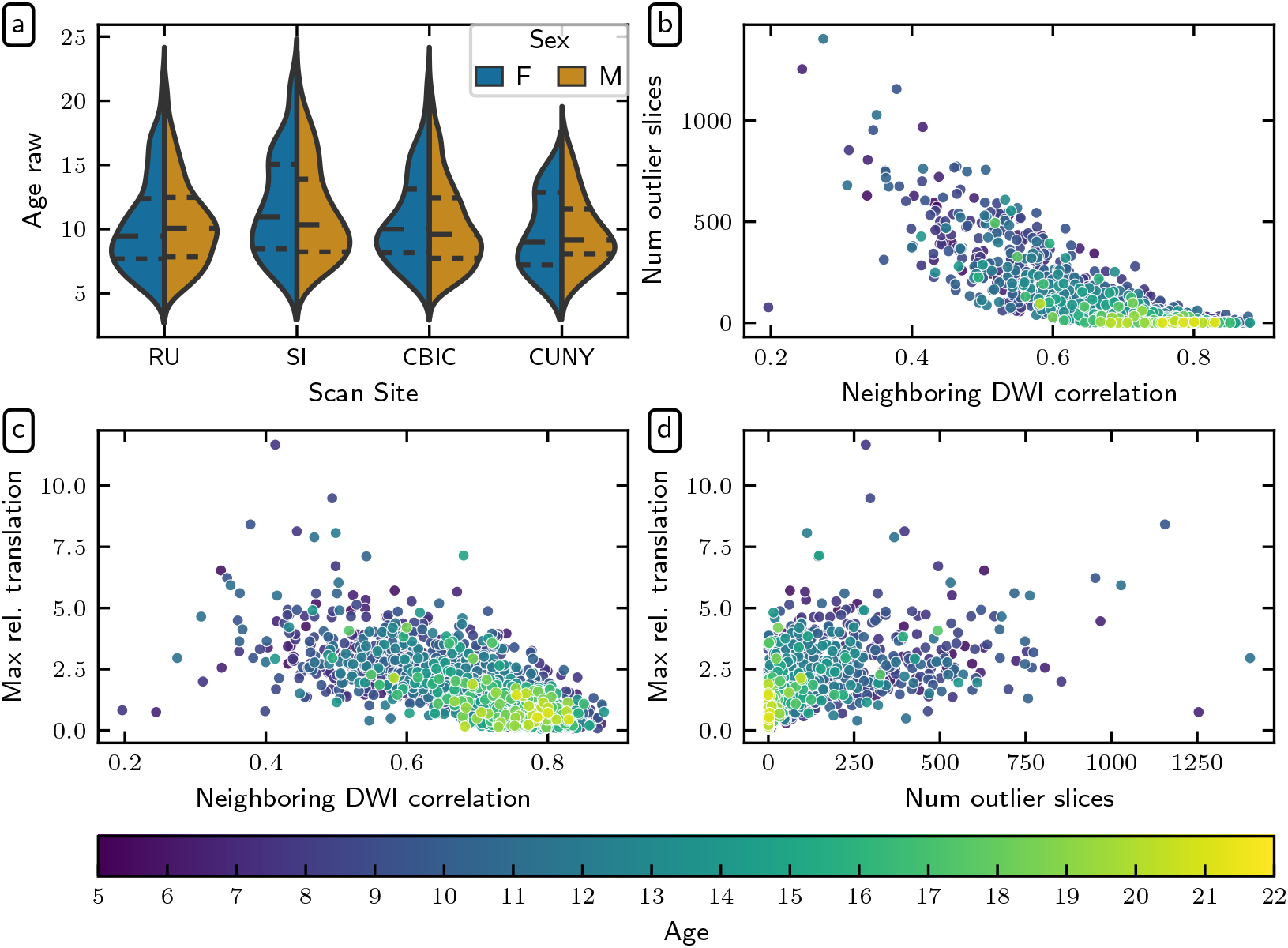
Demographic and *QSIPrep* quality metric distributions: **(a)** HBN age distributions by sex for each scanning site. Dashed lines indicate age quartiles. The remaining plots show associations between **(b)** neighboring diffusion-weighted imaging (DWI) correlation (***Yeh et al., 2019***) and the number of outlier slices, **(c)** neighboring DWI correlation and maximum relative translation, and **(d)** the number of outlier slices and maximum relative translation. The number of outlier slices is positively associated with the maximum relative translation, while neighboring DWI correlation is negatively associated with the other two metrics. These plots are colored by age, and reveal that older participants generally have higher quality data.

### Healthy Brain Network Preprocessed Open Diffusion Derivatives

Curated BIDS data and their corresponding *QSIPrep* outputs are provided in the FCP-INDI Amazon Web Services (AWS) S3 bucket ^2^. This public resource can be accessed by anyone using standard S3 access tools.

The curation process accounts for the acquisition variability inherent in large multi-site datasets by identifying unique *variants* in the HBN dMRI and eldmap acquisitions. Each session was grouped according to metadata parameters that affect the dMRI signal (PhaseEncodingDirection, EchoTime, VoxelSize, FlipAngle, PhasePartialFourier, NumberOfVolumes, Fieldmap availability). Using the “Curation of BIDS” (CuBIDS) package (***Covitz et al., 2022***), we identified a total of 20 unique DWI acquisitions across HBN-POD2, where about 5% of acquisitions were different from the most common DWI acquisition at their site. The specific variant of each session is provided as a column in the participant.tsv file and a summary of variants with participant counts is provided in Appendix 1.

The processed diffusion derivatives are standard *QSIPrep* outputs, which contain preprocessed imaging data along with the corresponding QC metrics:

#### • *Anatomical Data*

Preprocessed images, segmentations and transforms for spatial normalization are located in the anat/ directory of each session. The gray matter, white matter and cerebrospinal fluid (`GM, WM, CSF`) probabilistic segmentations are provided in nifti format with the `_probtissue` suffix. The deterministic segmentation is in `_dseg.nii.gz`. All images are in alignment with AC-PC-aligned `sub-X_desc-preproc_T1w.nii.gz` image unless they have `space-MNI152NLin2009cAsym` in their file name, in which case they are aligned to the MNI Nonlinear T1-weighted asymmetric brain template (version 2009c; (***Fonov et al., 2009a***)). The spatial transform between the AC-PC T1w image and the MNI space brain is in the ITK/ANTs format file named `sub-X_from-MNI152NLin2009cAsym_to-T1w_mode-image_xfm.h5`. The brain mask from `ANTsBrainExtraction.sh` is included in the file with the `_desc-brain_mask.nii.gz` suffix.

#### • *Diffusion Data*

The preprocessed dMRI scan and accompanying metadata are located in the `dwi/` directory of each session. The fully-preprocessed dMRI data is named according to the file pattern `sub-X_space-T1w_desc-preproc_dwi.nii.gz`. These images all have an isotropic voxel size of 1.7 mm and are aligned in world coordinates with the anatomical image located at `anat/sub-X_desc-preproc_T1w.nii.gz`. Gradient information is provided in `bval/bvec` format compatible with DIPY and DSI Studio and the `.b` format compatible with MRtrix3. Volume-wise QC metrics including head motion parameters are included in the confounds. `tsv` file. Automatically computed quality measures for the entire image series are provided in the `ImageQC.csv` file, which includes the neighboring DWI Correlation, number of bad slices and head motion summary statistics. Figure 2 depicts pairwise distributions for the three of these automated data quality metrics that were most informative in QC models described later (see Appendix 3 for further details). The `desc-brain_mask` file is a dMRI-based brain mask that should only be used when the T1w-based brain mask is inappropriate (i.e. when no susceptibility distortion correction has been applied).

### Quality Control

To QC all available HBN dMRI data, we adopted a hybrid QC approach that combines expert rating, community science, and deep learning, drawing on the success of a previous application in assessing the quality of HBN’s structural T1w MRI data (***Keshavan et al., 2019***). This method (i) starts with dMRI expert raters labelling a small subset of participants, the “gold standard” dataset; (ii) ampli es these labels using a community science web application to extend expert ratings to a much larger subset of the data, the community science subset and (iii) trains a deep learning model on the community science subset to predict expert decisions on the entire dataset.

#### Expert quality control

To create a gold standard QC dataset, we first developed *dmriprep-viewer*, a dMRI data viewer and QC rating web application to display *QSIPrep* outputs and collect expert ratings (***Richie-Halford et al., 2022***). Six of the co-authors, who are all dMRI experts, rated a 200-participant subset of the HBN-POD2 data using extensive visual examination of each participant’s dMRI data, including the preprocessed diffusion weighting imaging (DWI) time series, a plot of motion parameters throughout the DWI scan, and full 3D volumes depicting (i) the brain mask and *b* = 0 to T1w registration and (ii) a directionally encoded color fractional anisotropy (DEC-FA) image laid over the *b* = 0 volume. See Appendix 2 for an example of the *dmriprep-viewer* interface. The experts rated participants using a ve-point scale with ratings of “definitely fail,” “probably fail,” “not sure,” “probably pass,” and “definitely pass.” The distribution of scores given by the experts demonstrates that the gold standard dataset included a range of data quality (Figure 3a). Mean expert ratings correlated with the three *QSIPrep* automated QC metrics that were most informative for the XGB model described in the next section: neighboring DWI correlation ***(Yeh et al., 2019)*** (Figure 3b), maximum relative translation (Figure 3c), and number of outlier slices (Figure 3d). The neighboring DWI correlation characterizes the pairwise spatial correlation between pairs of DWI volumes that sample neighboring points in *q*-space. Since lower values indicate reduced data quality, it is reassuring that the neighboring DWI correlation correlated directly with expert ratings (Pearson CC: 0.797). Conversely, high relative translation and a high number of motion outlier slices reflect poor data quality and these metrics were inversely related to mean expert rating (Pearson CC: −0.692 and Pearson CC: −0.695, respectively).

**Figure 3.**
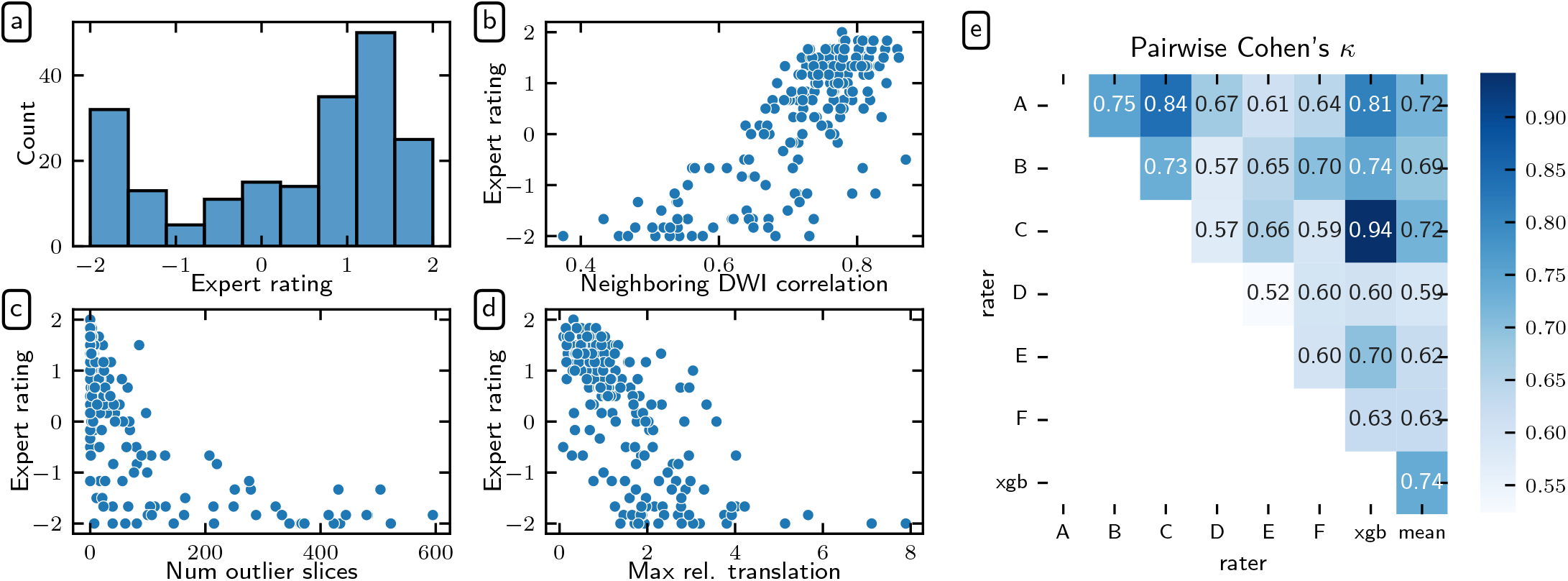
Expert QC results: Six dMRI experts rated a subset of 200 participants. Experts agreed with *QSIPrep*’s automated QC metrics. Here we show the distribution of mean expert QC ratings **(a)** and associations between the mean expert QC rating and the *QSIPrep* metrics **(b)** neighboring diffusion-weighted imaging (DWI) correlation (***Yeh et al., 2019***), **(c)** maximum relative translation, and **(d)** number of outlier slices. As expected, neighboring DWI correlation is directly correlated with expert rating while the other two metrics are inversely correlated with expert rating. **(e)** Experts agreed with each other. Here we show the pairwise Cohen’s *κ* measure of inter-rater reliability (see text for ICC calculations). The XGB model has an inter-rater reliability (quantified here as Cohen’s *κ*) that is indistinguishable from the other raters

In addition to agreeing qualitatively with *QSIPrep*’s automated QC metrics on average, the expert raters also tended to agree with each other (Figure 3e). We assessed inter-rater reliability (IRR) using the pairwise Cohen’s *κ* ***(Di Eugenio and Glass, 2004***), which exceeded 0.52 in all cases, and with a mean value of 0.648. In addition to the pairwise Cohen’s *κ*, we also computed the intra-class correlation (ICC) ***(Hallgren, 2012)*** as a measure of IRR. ICC3k is the appropriate variant of the ICC to use when a fixed set of *k* raters each code an identical set of participants, as is the case here. ICC3k for inter-rater reliability among the experts was 0.930 (95% CI: [0.91, 0.94]), which is qualitatively considered an “excellent” level of IRR ***(Cicchetti, 1994***). The high IRR provides confidence that the average of the expert ratings for each image in the gold standard is an appropriate target to use for training a machine learning model that predicts the expert scores.

#### Community science quality control

Although the expert raters achieved high IRR and yielded intuitive associations with *QSIPrep*’s automated QC metrics, generating expert QC labels for the entire HBN-POD2 dataset would be prohibitively time consuming. To assess the image quality of the remaining participants, we deployed *Fibr* (https://fibr.dev), a community science web application in which users assigned binary pass/fail labels assessing the quality of horizontal slice DEC-FA images overlaid on the *b* = 0 image (see Appendix 2 for an example). specifically, *Fibr* users saw individual slices or an animated sequence of ten slices taken from the entire DEC-FA volume that the expert raters saw. The *Fibr* users, therefore, saw only a subset of the imaging data that the dMRI experts had access to for a given participant, but they saw data from many more participants. In total, 374 community scientists provided 587,778 ratings for a mean of > 50 ratings per slice (or > 200 ratings per participant) from 1653 participants. Of the community scientists, 145 raters provided > 3,000 ratings each and are included in the *Fibr* Community Science Consortium as co-authors on this paper ***(Ward-Fear et al., 2020)*** (see Appendix 4 for a list of consortium co-authors).

There are three issues to account for when comparing *Fibr* and expert QC ratings. First, the unadjusted *Fibr* ratings were overly optimistic; i.e., on average, community scientists were not as conservative as the expert raters (Figure 4a). Second, different community scientists provide data of differing accuracy. That is, they were less consistent across different views of the same image, and/or were less consistent with expert ratings for the same data). This means that data from some *Fibr* raters was more informative than others. Third, important information about data quality was provided in the *QSIPrep* data quality metrics, which were not available to *Fibr* raters. To account for rater variability and take advantage of the information provided by *QSIPrep*, we trained gradient boosted decision trees ***(Chen and Guestrin, 2016a)*** to predict expert scores, scaled to the range [0, 1] and binarized with a 0.5 threshold, based on a combination of community science ratings and automated *QSIPrep* QC metrics. One can think of the gradient boosting model as assigning more weight to *Fibr* raters who reliably agree with the expert raters, thereby resolving the aforesaid issues with community rater accuracy. We refer to this gradient boosting model as XGB.

**Figure 4.**
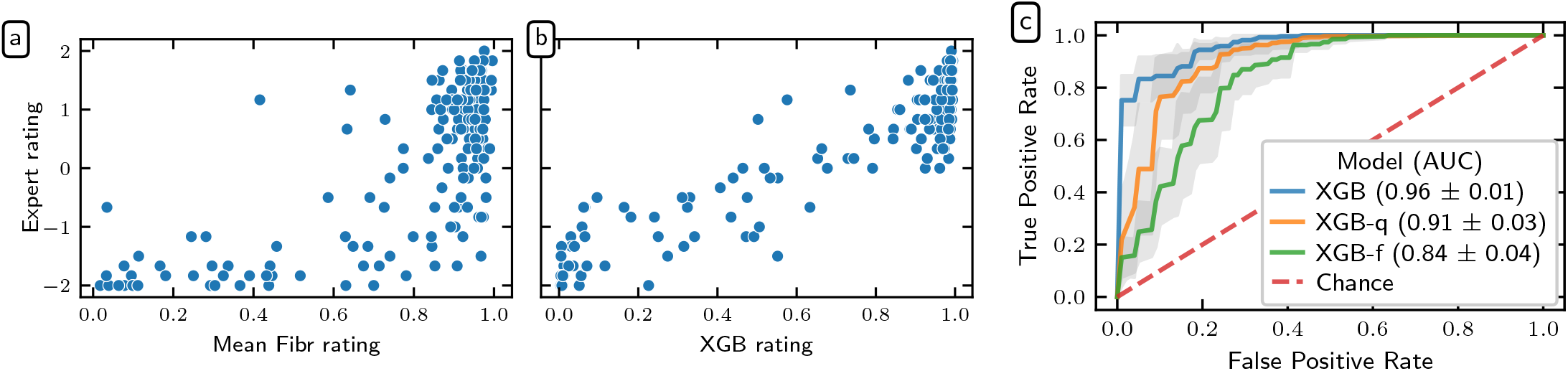
Community science predictions of the expert ratings: Scatter plots showing the relationship between mean expert rating and both mean *Fibr* rating **(a)** and XGB prediction **(b)**. *Fibr* raters overestimated the quality of images compared to expert raters. But the XGB prediction compensated for this by incorporating automated QC metrics and weighting more valuable *Fibr* raters. **(c)** ROC curves for the XGB, XGB-q, and XGB-f models. Translucent bands represent one standard deviation from the mean of the cross-validation splits.

To clarify the contributions of the automated QC metrics and the community science raters, we trained two additional gradient boosting models: (i) one trained only on the automated *QSIPrep* data quality metrics, which we call XGB-q and (ii) one trained on only the *Fibr* ratings, which we call XGB-f. XGB-f may be viewed as a data-driven weighting of community scientists’ ratings, while XGB-q may be viewed as a generalization of data quality metric exclusion criteria. XGB, combining information from both *Fibr* ratings and *QSIPrep* data quality metrics attained a cross-validated area under the receiver operating curve (ROC-AUC) of 0.96 ± 0.01 on the “gold standard,” where the ± indicates the standard deviation of scores from repeated *k*-fold cross-validation (Figure 4b). In contrast, XGB-q attained an ROC-AUC of 0.91 ± 0.03 and XGB-f achieved an ROC-AUC of 0.84 ± 0.04. The enhanced performance of XGB-q over XGB-f shows that community scientists alone are not as accurate as automated data quality metrics are at predicting expert ratings. And yet, the increased performance of XGB over XGB-q demonstrates that there is additional image quality information to be gained by incorporating community scientist input.

As a way of evaluating the quality of the XGB predictions, consider the fact that the average Cohen’s *κ* between XGB and the expert raters was 0.74, which is higher than the average Cohen’s *κ* between any of the other raters and their human peers (Figure 3). This is not surprising, given that the XGB model was fit to optimize this match, but further demonstrates the goodness of fit of this model.

Nevertheless, this provides confidence in using the XGB scores in the next step of analysis, where we treat the XGB model as an additional coder and extend XGB ratings to participants without *Fibr* ratings. In this case, when a subset of participants is coded by multiple raters and the reliability of their ratings is meant to generalize to other participants rated by only one coder, the single-measure ICC3, as opposed to ICC3k, should be used. When adding XGB to the existing expert raters as a seventh expert, we achieved **ICC3** = 0.709(95%*CI*: [0.66, 0.75]). The high ICC3 value after inclusion of the XGB model justifies using the XGB scores as the target for training an image-based deep learning network.

#### Automated quality control labelling through deep learning

While the XGB “rater” does a good job of extending QC ratings to the entire community science subset, this approach requires *Fibr* scores; without community science *Fibr* scores, only the less accurate XGB-q prediction can be employed. Consequently, a new, fully automated QC approach is needed that can be readily applied to new data releases from HBN.

We therefore trained a deep convolutional neural network to predict the XGB ratings directly from *QSIPrep* outputs. We modified an existing 3D convolutional neural network (CNN) architecture (***Zunair et al., 2020***)—previously applied to the ImageCLEF Tuberculosis Severity Assessment 2019 benchmark (***Dicente Cid et al., 2019***)—to accept multichannel input generated from the preprocessed dMRI: the *b* = 0 reference diffusion image, each of the three cardinal axis components of the DEC-FA image, and, optionally, automated QC metrics from *QSIPrep*. We trained this network on XGB scores and validated it against the gold standard expert-scored dataset. We refer to the convolutional neural network model trained only on imaging data as CNN-i and the model that incorporates automated QC metrics as CNN-i+q. The two models performed nearly identically and achieved an ROC-AUC of 0.947 ± 0.004 (Figure 5a). The near-identical performance suggests that *QSIPrep*’s automated data quality metrics provided information that was redundant with information available in the imaging data. Both CNN-i and CNN-i+q outperformed XGB-q, which was trained only on automated QC metrics, but both modestly underperformed relative to the full XGB model, that uses *Fibr* scores in addition to the *QSIPrep* data quality metrics.

**Figure 5.**
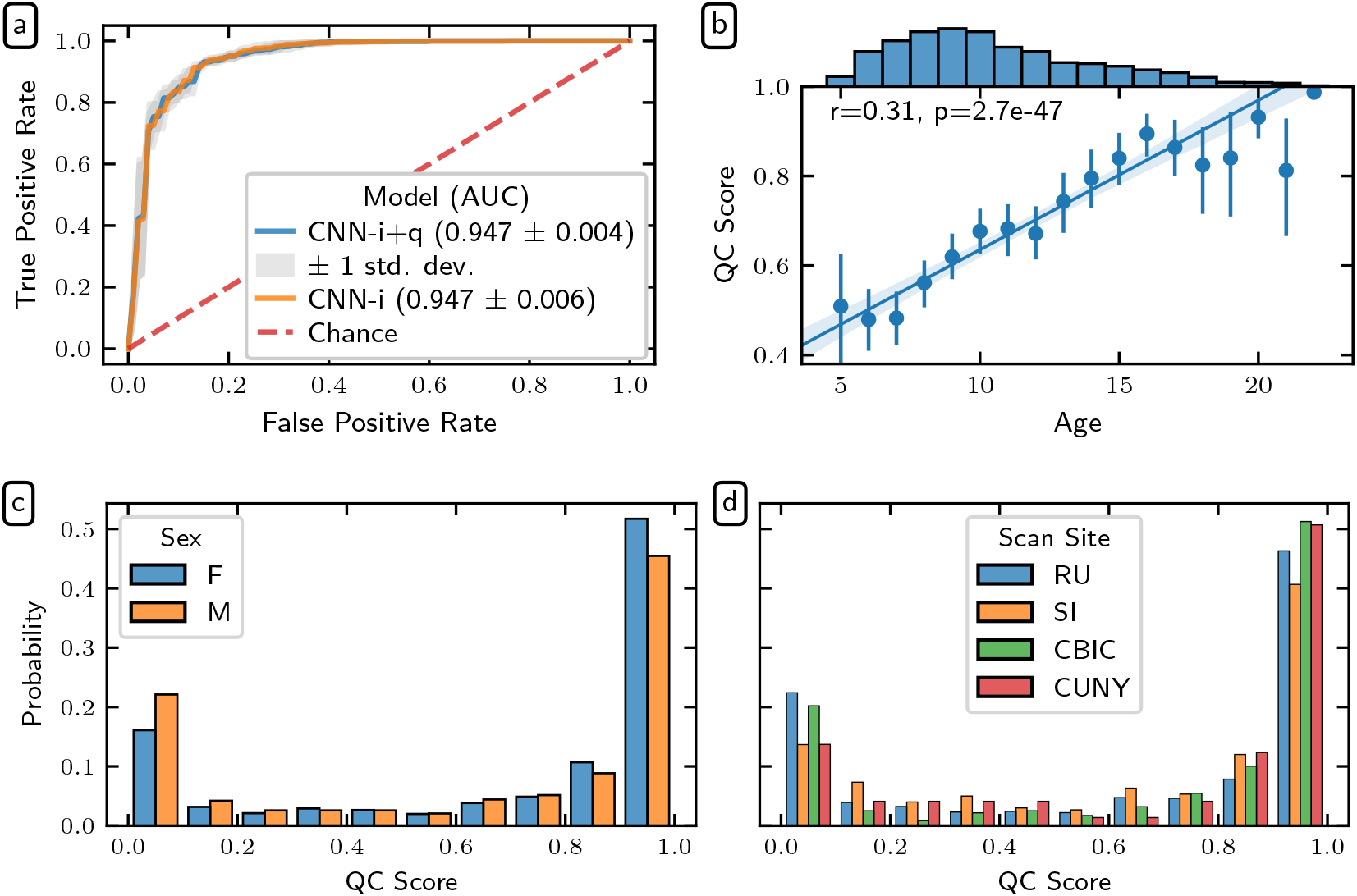
Deep learning QC scores: **(a)** ROC curves for two deep learning models trained on imaging data: one trained with additional automated data quality metrics from *QSIPrep* (blue) and one trained without (orange). The models performed roughly identically, reflecting that the data quality metrics are derived from the imaging data and are therefore redundant. Both outperformed the XGB-q predictions, indicating the added value of the diffusion weighted images. However, both models underperformed the XGB predictions, which also incorporate information from *Fibr* ratings for each scan. The error bands represent one standard deviation from the mean of the cross-validation splits. **(b)** Joint distributions showing a strong direct association between age and QC score (Pearson CC: 0.31). This likely reflects the well-known negative association between age and head motion in pediatric neuroimaging. The dots encode the mean QC score for each year of age with error bands representing a bootstrapped 95% confidence interval. The line depicts a linear regression relating age and QC score with translucent bands encoding a bootstrapped 95% confidence interval. Histograms showing the relationship between participants QC scores and their sex **(c)** and scan site **(d)**. QC distributions are independent of sex and scanning site.

The openly available HBN-POD2 data released with this paper provides four QC ratings: the mean expert QC ratings, XGB-q and XGB predicted scores, as well as the CNN-i predicted score. However, we treat the CNN-i score as the definitive QC score because it is available for all participants, can be easily calculated for new participants in future HBN releases, and is more accurate than XGB-q in predicting expert ratings in the “gold standard” report set. When we refer to a participant’s QC score without specifying a generating model, the CNN-i score is assumed. Figure 5 depicts the distribution of these QC scores by age (Figure 5b), sex (Figure 5c), and scanning site (Figure 5d). QC distributions are similar for each scan site and for male and female participants ^3^.

#### Attribution masks for the deep learning classifier

We generated post-hoc attribution maps that highlight regions of the input volume that are relevant for the QC score. The integrated gradient method (***Sundararajan et al., 2017***) is a gradient-based attribution method (***Ancona et al., 2019***) that aggregates gradients for synthetic images interpolating between a baseline image and the input image. It has been used to interpret deep learning models applied to retinal imaging in diabetic retinopathy (***Sayres et al., 2019***) and glaucoma (***Mehta et al., 2021***) prediction, as well as in multiple sclerosis prediction from brain MRI (***Wargnier-Dauchelle et al., 2021***). Our goal is to confirm that the CNN-i model was driven by the same features that would drive the expert rating, thereby bolstering the decision to apply it to new data.

Figure 6 shows attribution maps for example participants from each confusion class: true positive, true negative, false positive, and false negative. The columns correspond to the different channels of the deep learning input volume: the *b* = 0 reference image and the DEC-FA in the *x, y*, and *z* directions. The blue voxels indicate positive attribution, that is, data that supports a passing QC classification. Conversely, the red voxels indicate negative attribution, data that supports a failing QC classification. The true positive map indicates that the network was looking at the entire brain rather than focusing on any one anatomical region (Figure 6a). Moreover, the model identified white matter fascicles that travel along the direction of the input channel: lateral for *x*, anterior-posterior for *y*, and superior-inferior for *z*. The true negative attribution map (Figure 6b) reveals that when the reference *b* = 0 volume contains motion artifacts, such as banding, the network ignored the otherwise positive attributions for the clearly identifiable white matter tracts in the DEC-FA channels. The false positive map (Figure 6c) and the false negative map (Figure 6d) should be interpreted differently since they come from low confidence predictions; the probability of passing hovered on either side of the pass/fail threshold. For example, in the false positive case, the network was confused enough that it treated voxels that are outside of the brain to be as informative as voxels in the major white matter bundles.

**Figure 6.**
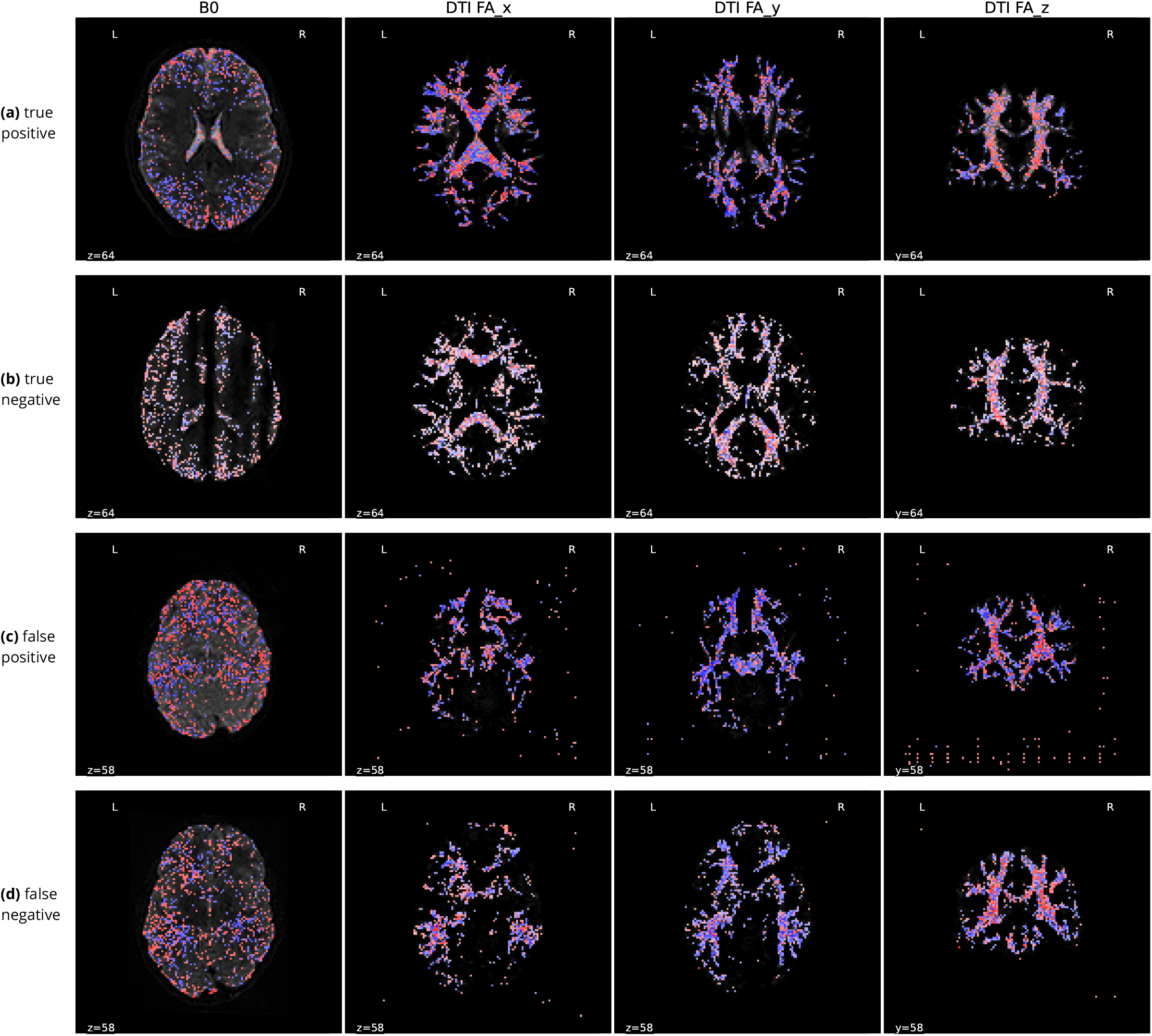
Integrated gradients attribution maps for the deep learning classifier: Each column depicts a different channel of the input tensor: the *b* = 0 DWI volume and the DEC-FA images in the *x, y*, and *z* directions. The first three columns show an axial slice while the last column shows a coronal slice. Blue voxels indicate positive attribution (i.e. evidence for passing the participant), while red voxels indicate negative attribution (i.e. evidence for QC failure). The underlying grayscale depicts the input channel. Each row depicts a representative participant from each confusion class: **(a)** Attribution maps for a true positive prediction. The model looked at the entire brain and focused on known white matter bundles in the DEC-FA channels. In particular, it focused on lateral bundles in the *x* direction, anterior-posterior bundles in the *y* direction, and superior-inferior bundles in the *z* direction. **(b)** Attribution maps for a true negative prediction. The model focused primarily on the *b* = 0 channel, suggesting that it ignores DEC-FA when motion artifacts like banding are present. **(c)** Attribution maps for a false positive prediction. Both the false positive and negative predictions were low confidence predictions. This is reinforced by the fact that the model viewed some voxels that are outside of the brain as just as informative as those in major white matter tracts. **(d)** Attribution maps for a false negative prediction. The model failed to find long-range white matter tracts in the anterior-posterior and lateral directions. We also speculate that the model expected left-right symmetry in the DEC-FA channels and assigned negative attribution to asymmetrical features.

#### QC prediction models can generalize to unseen sites

Site harmonization is a major issue for any multisite neuroimaging study and developing automated QC tools that generalize between sites has been a perennial issue (***Esteban et al., 2017***). Furthermore, the ability to generalize between sites in a single multisite study would signal the promise of generalizing to other datasets altogether. To better understand the ability of our QC models to generalize across scanning sites, we trained several variants of the XGB-q and CNN-i models on partitions of the data with different sites held out (Figure 7). ROC-AUC for generalization is uniformly high for both the XGB-q and the CNN-i models (Table 1). However, more importantly, accuracy and balanced accuracy vary substantially: depending on the site that was used for training, balanced accuracy could be as low as guess rate, particularly for the CNN-i model. Notably, it seems that including the RU site in the training data led to relatively high balanced accuracy in both models. The XGB-q model balanced accuracy was less dependent on the specific sites used for training, but also displayed some variability across permutations of this experiment. In particular, the benefit from including the “right site” in the training data, namely RU, eclipsed the slight benefit conferred by including more than one site in the training data.

**Table 1.** Site generalization summary statistics: Below we list the mean ± standard deviation of the site generalization evaluation metrics displayed in Figure 7. For each of the CNN-i and XGB-q model families and each of the site generalization splits, we report the accuracy, balanced accuracy, and ROC-AUC.

| Model | Site | Accuracy | Balanced accuracy | ROC-AUC |
| --- | --- | --- | --- | --- |
| CNN-i | train: CBIC + CUNY, test: RU | $0.748 \pm 0.086$ | $0.652 \pm 0.112$ | $0.930 \pm 0.015$ |
| | train: CBIC, test: RU + CUNY | $0.696 \pm 0.095$ | $0.574 \pm 0.123$ | $0.791 \pm 0.169$ |
| | train: RU + CUNY, test: CBIC | $0.859 \pm 0.033$ | $0.847 \pm 0.030$ | $0.912 \pm 0.013$ |
| | train: RU, test: CBIC + CUNY | $0.851 \pm 0.018$ | $0.753 \pm 0.029$ | $0.910 \pm 0.014$ |
| XGB-q | train: CBIC+CUNY, test: RU | $0.763 \pm 0.071$ | $0.805 \pm 0.052$ | $0.895 \pm 0.006$ |
| | train: CBIC, test: RU+CUNY | $0.725 \pm 0.079$ | $0.779 \pm 0.058$ | $0.886 \pm 0.019$ |
| | train: RU+CUNY, test: CBIC | $0.894 \pm 0.024$ | $0.838 \pm 0.036$ | $0.931 \pm 0.018$ |
| | train: RU, test: CBIC+CUNY | $0.886 \pm 0.030$ | $0.816 \pm 0.048$ | $0.940 \pm 0.017$ |

**Figure 7.**
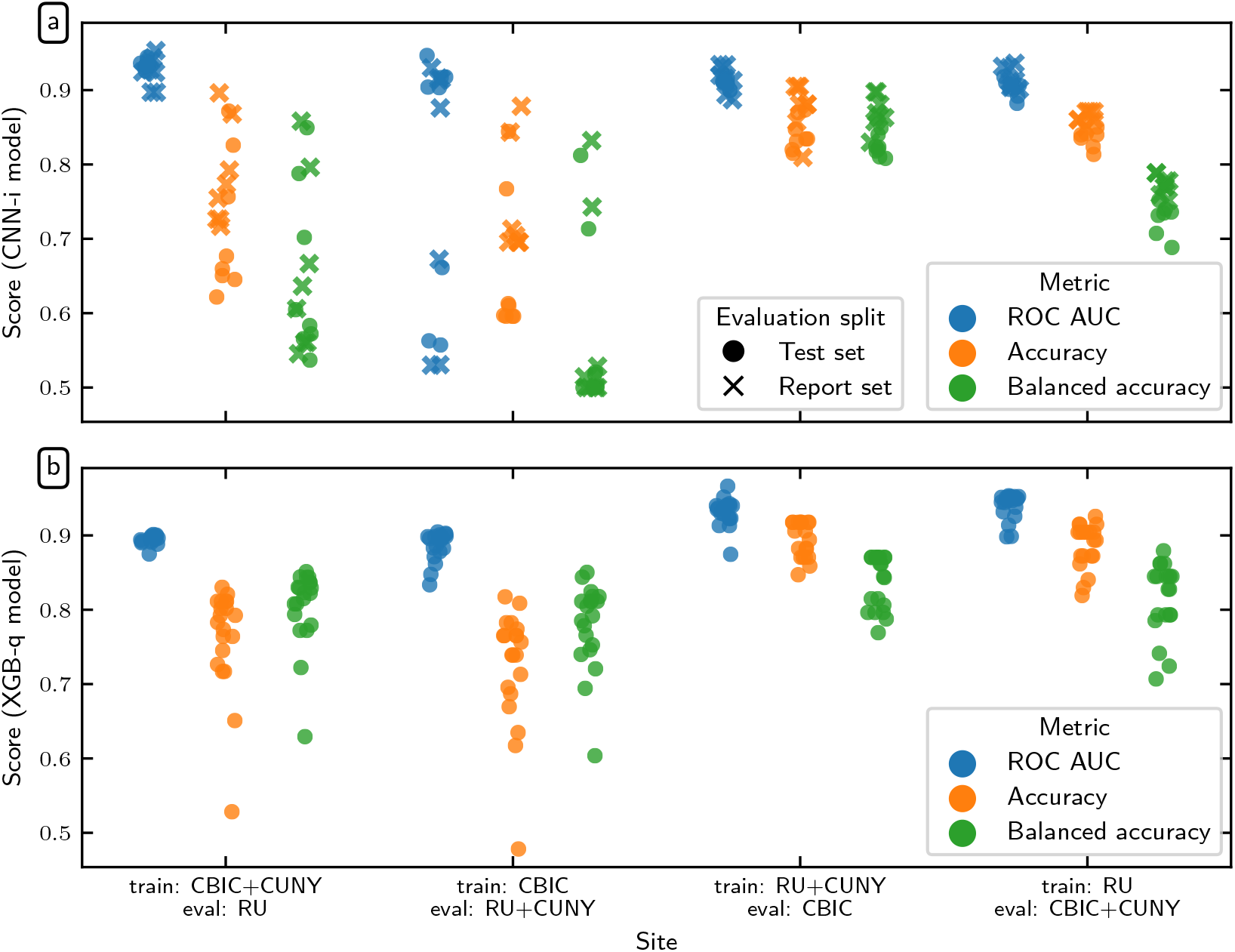
Generalization of QC scores to unseen sites: In each experiment, CNN-i (**a**) and XGB-q (**b**) models were trained with some sites held out and evaluated only on data from these held out sites. Model performance is quantified as ROC-AUC (blue), accuracy (orange) and balanced accuracy (green). For XGB-q, the targets are the expert ratings on data from the held out site. For CNN-i, performance is scored against XGB scores (as used before; test set in lled circles), or expert ratings on the data from the held out site (report set in crosses). Summary statistics for this plot are listed in Table 1.

### Quality control improves inference

To demonstrate the effect that quality control has on inference, we analyzed tract profile data derived from HBN-POD2 data. Tract profiling (***Yeatman et al., 2012***; ***Jones et al., 2005***; ***Colby et al., 2012***; ***O’Donnell et al., 2009***; ***Kruper et al., 2021***) is a subset of tractometry (***Jones et al., 2005***; ***Bells et al., 2011***), which uses the results of dMRI tractography to quantify properties of the white matter along major pathways. Tract-profiling retains the values of diffusion metrics along the trajectory of each bundle of tractography streamlines, rather than computing summary statistics summarized at the level of each bundle. In Figure 8, we plot mean diffusivity tract profiles grouped into four QC bins along the length of twenty-four bundles: While some bundles, such as the cingulum cingulate (CGC) and the inferior longitudinal fasciculus (ILF), appear insensitive to QC score, others, such as the uncinate (UNC) and the orbital portion of the corpus callosum, exhibit strong differences between QC bins. In most bundles, low QC scores tend to flatten the profile, indicating that mean diffusivity appears artifactually homogeneous across the bundle.

**Figure 8.**
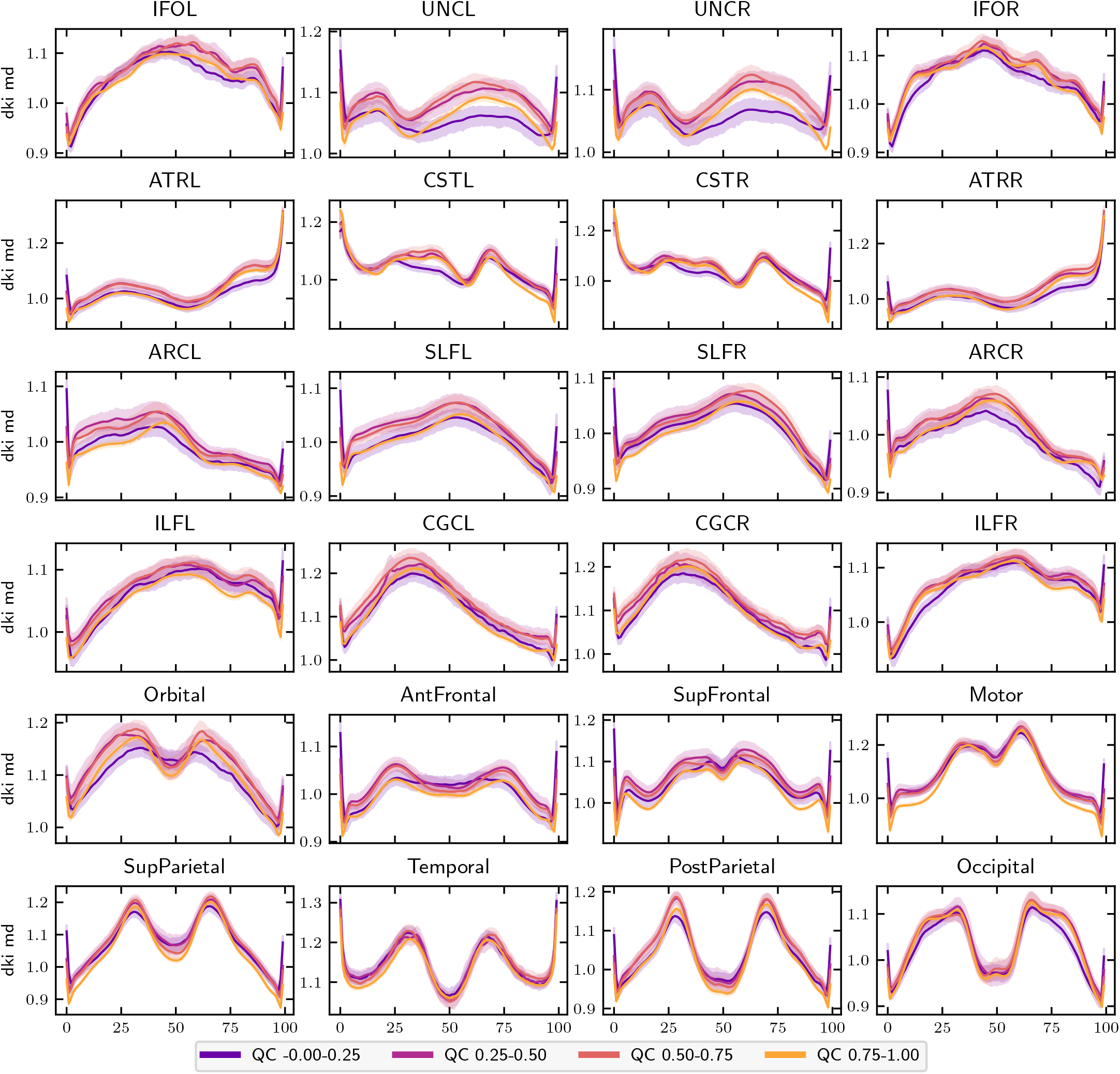
MD bundle profiles show large QC group differences: MD profiles binned by QC score in twenty-four major while matter bundles. The *x*-axis represents distance along the length of the fiber bundle. The left and right uncinate bundles were the most sensitive to QC score. Generally, QC score tended to flatten bundle profiles. Error bands represent bootstrapped 95% confidence intervals. Bundle abbreviations for lateralized bundles contain a trailing “L” or “R” indicating the hemisphere. Bundle abbreviations: inferior fronto-occipital fasciculus (IFO), uncinate (UNC), thalamic radiation (ATR), corticospinal (CST), arcuate (ARC), superior longitudinal fasciculus (SLF). inferior longitudinal fasciculus (ILF), cingulum cingulate (CGC), orbital corpus callosum (Orbital), anterior frontal corpus callosum (AntFrontal), superior frontal corpus callosum (SupFrontal), motor corpus callosum (Motor), superior parietal corpus callosum (SupParietal), temporal corpus callosum (Temporal), post-parietal corpus callosum (PostParietal), and occipital corpus callosum (Occipital). **Figure 8–Figure supplement 1.** FA bundle profiles

The effect of QC score on white matter bundle profiles indicates that researchers using HBN-POD2 should incorporate QC in their analyses, either by applying a QC cut-off when selecting participants or by explicitly adding QC score to their inferential models. Failure to do so may cause spurious associations or degrade predictive performance. To demonstrate this, we selected participant age as a representative phenotypic benchmark because (i) it operates on a natural scale with meaningful units and (ii) despite the unique methodological challenges it presents for biomarker identi cation (***Nelson et al., 2020***), brain age prediction may be diagnostic of overall brain health (***Franke et al., 2010***; ***Cole et al., 2019***; ***Richie-Halford et al., 2021***). We observed the effect of varying QC cutoff on the predictive performance of an age prediction model. Cross-validated 2 scores for an age prediction model varied depending on the QC cut off (Figure 9). An initial large improvement was achieved by excluding the 200 participants with the lowest QC scores, followed by a gradual increase in performance. Finally, when a large number of participants is excluded, performance deteriorated again.

**Figure 9.**
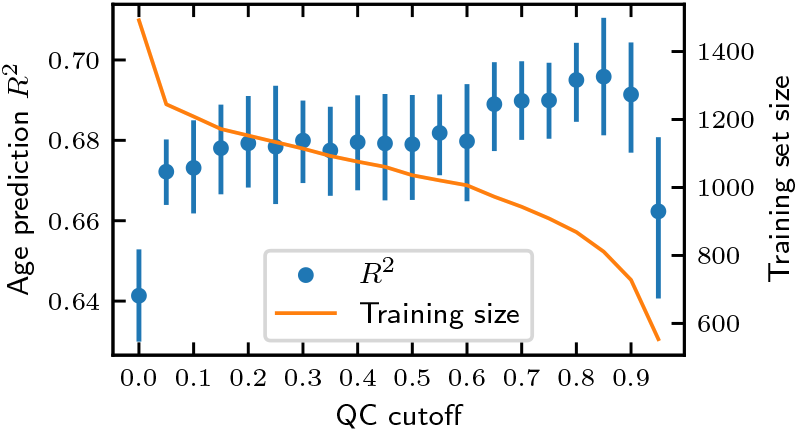
Imposing a QC cutoff improves age prediction: Cross validated *R*^2^ scores (left axis, blue dots) from an age prediction model increase after screening participants by QC score. We see the most dramatic increase in *R*^2^ after imposing even the lowest cutoff of 0.05. Thereafter, the *R*^2^ scores trend upward until a cutoff of ~0.95, where the training set size (right axis, orange line) becomes too small to sustain model performance. The error bands represent a bootstrapped 95% confidence interval.

## Discussion

We present HBN-POD2, one of the largest child and adolescent diffusion imaging datasets with preprocessed derivatives that is currently openly available. The dataset was designed to comply with the best practices of the field. For example, it complies with the current draft of the BIDS diffusion derivative Specification (***Pestilli et al., 2021***). It will grow continuously as the HBN study acquires more data, eventually reaching its 10,000 participant goal.

### Preprocessing and quality control increase the impact of openly-available data

The most immediate contribution of this work is a large analysis-ready dMRI data resource, openly accessible to the public. In the past decade, projects such as the Human Connectome Project (HCP) (***Van Essen et al., 2013***), UK Biobank (***Miller et al., 2016***), ABCD (***Jernigan and Brown, 2018***), and CamCAN (***Taylor et al., 2017***; ***Shafto et al., 2014***) and of course FCP-INDI (which includes HBN) (***Mennes et al., 2013***) have ushered a culture of data sharing in open big-data neuroscience. The adoption and reuse of these datasets reduces or eliminates the data collection burden on downstream researchers. Some projects, such as the HCP (***Glasser et al., 2013***), also provide preprocessed derivatives, further reducing researchers’ burden and extending the benefits of data-sharing from data collection to preprocessing and secondary analysis. Following the example of the HCP, HBN-POD2 provides analysis-ready dMRI derivatives. This avoids duplication of and heterogeneity across the preprocessing effort while also ensuring a minimum standard of data quality for HBN researchers. We also provide the CuBIDS variant annotation in the participants.tsv file, allowing researchers to account for the imaging heterogeneity inherent in a dataset of this size. Making MRI derivatives accessible not only reduces the burden of processing large datasets for research groups with limited resources (***Laird, 2021***), but also aids research performed by clinicians who are interested in brain-behavior relationships but may be lacking the technical training to process large-scale dMRI data.

The data is amenable to many different analyses, including tractometry (***Yeatman et al., 2012***, ***2018***; ***Kruper et al., 2021***), graph theoretical analysis (***Yeh et al., 2020***), and combinations with functional MRI data and other data types for the same participants. The availability of standardized preprocessed diffusion data will allow researchers to create and test hypotheses on the white matter properties underlying behavior and disease, from reading and math acquisition to childhood adversity and mental health. As such, this dataset will accelerate discovery at the nexus of white matter microstructure and neurodevelopmental and learning disorders.

In large developmental datasets, it is critically important to perform accurate and reliable QC of the data. QC is associated not just with age, but with many phenotypic variables of interest in cognition and psychopathology (***Siegel et al., 2017***). HBN-POD2 provides four separate QC scores alongside its large dataset of pediatric neuroimaging diffusion derivatives, paving the way for users of the data to incorporate considerations of data quality into their analysis of the processed data. Unsurprisingly, QC scores are strongly correlated with age (Figure 5). This accords with the negative association between head motion and age in developmental studies, which is well established both in general (***Power et al., 2012***; ***Satterthwaite et al., 2012***; ***Fair et al., 2012***; ***Yendiki et al., 2014***) and specifically for resting-state fMRI in the HBN dataset (***Alexander et al., 2017***). Moreover, it is important that QC has bundle-specific and spatially localized effects (Figure 8). Analysis of this data that does not incorporate QC is likely to find replicable but invalid effects. For example, in patient-control studies, patients are likely to have lower quality data. And analysis of such patient data that does not control for QC will find spatially-localized and replicable group differences that are due to data quality, not necessarily underlying neuroanatomical differences.

We further demonstrated the impact of QC in a benchmark age prediction task (Figure 9). In this case, the increase in model performance from imposing a QC cutoff is intuitive: we know from Figure 8 that participants with low QC scores have reduced MD, but MD also decreases as participants mature (***Yeatman et al., 2014***; ***Richie-Halford et al., 2021***). Eliminating participants with low QC therefore removes the ones who may look artificially older from the analysis, improving overall performance. The most noticeable improvement in performance comes after imposing the most modest cutoff of 0.05, suggesting that inferences may benefit from *any* QC screening. On the other hand, QC screening inherently introduces a tradeoff between the desire for high quality data and the desire for a large sample size. In this case, after a QC cutoff of around 0.9, the training set size is reduced such that it degrades predictive performance. Importantly, we do not expect the sensitivity analysis of an age prediction model to generalize to other analyses and therefore recommend that researchers using HBN-POD2 choose the most appropriate QC cutoff for their research question and consider including QC score as a model covariate in their analyses.

### Automated quality control: scalability, interpretability, and generalization

The predictive performance of the CNN-i model (Figure 5a) gives us confidence that it could accurately classify unseen data from the same sites, justifying its extension to the entire HBN-POD2 dataset and to future releases of HBN. However, one limitation of this model is that it does not satisfactorily explain its decisions. As deep learning models have been increasingly applied to medical image analysis, there is an evolving interest in the interpretability of these models (***Salahuddin et al., 2022***; ***Lipton, 2017***; ***Zech et al., 2018***; ***Ghassemi et al., 2021***). While an exhaustive interpretation of deep learning QC models is beyond the scope of this work, we provided a preliminary qualitative interpretation of the CNN-i model (Figure 6) that demonstrates the intuitive nature of its decisions.

The accuracy in generalizing to unseen data from HBN also suggested the tantalizing possibility that the QC models would be able to generalize to similar data from other datasets. To assess this, we trained the models with unseen sites held out (Figure 7). Both the CNN-i model and the XGB-q model do sometimes generalize to data from unseen sites, suggesting that they would be able to generalize to some other datasets as well. However, they do not reliably generalize, implying that they should not currently be used in this way. Future work could build upon the work that we have done here to establish a procedure whereby the models that we fit in HBN would be applied to data from other studies, but comprehensive calibration and validation would have to be undertaken as part of this procedure.

We recognize that decisions about QC inclusion must balance accuracy, interpretability, generalization to new data, and scalability to ever larger datasets. We therefore provide three additional scores: (i) the mean expert QC score for the 200 participants in the gold standard dataset, (ii) the scores predicted by the XGB model, which outperformed all other models when evaluated against the gold standard ratings, but which are only available for participants that have community science scores, and (iii) the scores predicted by the XGB-q model, which underperformed the deep learning generated scores, but which rely only on the automated QC metrics output by *QSIPrep*. We view the XGB-q scores, which are available for all participants, as a more interpretable and scalable fallback because the XGB-q model ingests *QSIPrep* output without any further postprocessing. XGB-q also provides slightly more uniform performance in generalization to unseen HBN sites (Figure 7). Because the XGB-q model most readily generalizes to other *QSIPrep* outputs, we package it as an independent QC service in the QSIQC software package (***Richie-Halford and Rokem, 2022b***), available both as a docker image at ghcr.io/richford/qsiqc and as a Streamlit app at https://share.streamlit.io/richford/qsiqc/main/app.py. The decision to use a more interpretable but slightly less performant method of generating QC scores was also advocated by (***Tobe et al., 2021***), who noted that the Euler number of T1-weighed images (***Rosen et al., 2018***) in the NKI-Rockland dataset can reliably predict scores generated with *Braindr*, the community science application developed in our previous work (***Keshavan et al., 2019***).

We also note that the issue of algorithmic impact in choosing a QC method is not exclusive to the deep learning model. We have chosen models that most reliably reproduce the gold standard ratings, but a reliable algorithm might still negatively influence researcher’s decisions. For example, excluding participants by QC score could spur them to exclude populations deserving of study, as when QC score is highly correlated with age or socio-economic status. We therefore caution researchers to examine interactions between the QC scores we provide and their phenotype of interest.

More generally, QC in the dataset that we have produced is fundamentally anchored to the decisions made by the expert observers. While Cohen’s *κ* between some pairs of experts can be as low as 0.52, IRR quantified across all of the experts with ICC3k is excellent. Nevertheless, it is possible that improvements to the final QC scores could be obtained through improvements to IRR, or by designing a more extensive expert QC protocol. The tradeoff between more extensive QC for each participant and more superficial QC on more participants was not explored in this study, but could also be the target for future research.

### Transparent pipelines provide an extensible baseline for future methods

While the primary audience of HBN-POD2 is researchers in neurodevelopment who will use the dMRI derivatives in their studies, other researchers may use HBN-POD2 to develop new preprocessing algorithms or quality control methods. In this respect, HBN-POD2 follows ***Avesani et al. (2019)***, who recognized the diverse interests that different scientific communities have in reusing neuroimaging data and coined the term *data upcycling* to promote multiple-use data sharing for purposes secondary to those of the original project. Complementing the approach taken in Avesani et al.’s work, which provided dMRI from a small number of participants preprocessed with many pipelines, HBN-POD2 contains many participants, all processed with a single state of the art pipeline, *QSIPrep*. For researchers developing new preprocessing algorithms, HBN-POD2 provides a large, openly available baseline to which they can compare their results.

Similarly, neuroimaging QC methods developers will benefit from a large benchmark dataset of expert, community science, and automated QC ratings, with which to test new methods. Importantly, the architecture and parameters of the deep learning network used for QC are also provided as part of this work, allowing application of this network to future releases of HBN data, and allowing other researchers to build upon our efforts. Indeed, in this work, we have extended our previous work on what we now call “hybrid QC”. This approach, which we originally applied to the first two releases of the HBN T1-weighted data (***Keshavan et al., 2019***) (using the *Braindr* web app: https://braindr.us) was extended here in several respects. First, the *Braindr* study used a smaller dataset of approximately 700 participants, while we extended this approach to well over 2000 participants. Second, *Braindr* relied on approximately 80,000 ratings from 261 users. Here, we received more than 500,000 ratings from 374 community scientists. As our understanding of the role of community scientist contributions has evolved, we decided that we would include as collective co-authors community scientists who contributed more than 3000 ratings (***Ward-Fear et al., 2020***). Third, *Braindr* used data from only a single site. Here, multi-site data was used. This opens up multiple possibilities for deeper exploration of between-site quality differences, and also for harmonization of QC across sites, as we have attempted here. Last, the most challenging extension of hybrid QC from *Braindr* to this study entailed developing an approach that would encompass multivolume dMRI data. On the one hand, this meant that the task performed by the expert observers was more challenging, because it required examination of the full dMRI time-series for every scan. To wit, expert inter-rater reliability was considerably higher for the T1-weighted only data in ***Keshavan et al.*** (***2019***) than for the dMRI data used (Figure 3e). On the other hand, it also meant that the 4D data had to be summarized into 2D data to be displayed in the *Fibr* web application. This was achieved by summarizing the entire time-series as a DEC-FA + *b* = 0 image and presenting community scientists with animated sections of these images that showed how the data extended over several horizontal slices. In addition, the extension to 4D data required developing new deep learning architectures for analysis of 4D images, including upstream contributions to *Nobrainer*, a community-developed software library for deep learning in neuroimaging data (***Kaczmarzyk et al., 2021***). These extensions demonstrate that the hybrid QC approach generalizes very well to a variety of different circumstances. Future applications of this approach could generalize to functional MRI data, as well as other large datasets from other kinds of measurements and other research domains.

### Future work and open problems

The HBN study plans to acquire imaging data for over 5000 participants, necessitating future data releases. Since future releases of HBN will also require future releases of HBN-POD2, a plan for these is essential. This is a general issue affecting multi-year neuroimaging projects for which derivative data is being released before study completion. The use of *QSIPrep*, *cloudknot* and the containerization of the QC score assignment process facilitate running the exact pipeline described in this paper on newly released participants. However, this approach is somewhat unsatisfactory because it fails to anticipate improvements in preprocessing methodology. That is, what should we do when *QSIPrep* is inevitably updated between HBN releases? Enforce standardization by using an outdated pipeline or use state-of-the-art preprocessing at the expense of standardized processing between releases? Because the use of *cloudknot* and AWS Spot Instances renders preprocessing fast and relatively inexpensive, we propose a third way: if improvements to the preprocessing pipeline are available with a new HBN release, we plan to execute the improved pipeline on the entire HBN dataset, while preserving the previous baseline release in an archived BIDS derivative dataset.

Undertaking the processing and QC effort to generate HBN-POD2 required construction and deployment of substantial informatics infrastructure, including tools for cloud computing, web applications for expert annotation and for community science rating and analysis software. All of these tools are provided openly, so that this approach can be generalized even more widely in other projects and in other scientific fields.

## Methods and Materials

To facilitate replicability, Jupyter notebooks (***Kluyver et al., 2016***) and Dockerfiles (***Merkel, 2014***) necessary to reproduce the methods described herein are provided in the HBN-POD2 GitHub repository at https://github.com/richford/hbn-pod2-qc. The specific version of the repository used in this study is documented in ***Richie-Halford and Rokem*** (***2022a***). The `make or make help` commands will list the available commands and `make build` will build the requisite Docker images to analyze HBN-POD2 QC data. In order to separate data from analysis code (***Wilson et al., 2017***), we provide intermediate data necessary to analyze the QC results in an OSF (***Foster and Deardorff, 2017***) project (***Richie-Halford and Rokem, 2021***), which can be downloaded using the `make data` command in the root of the HBN-POD2 GitHub repository. Most of the code in this repository uses Pandas (***McKinney, 2010***; ***pandas development team, 2020***), Numpy (***Harris et al., 2020***), Matplotlib (***Hunter, 2007***), and Seaborn (***Waskom, 2021***).

### Inputs

Inputs for this study consisted of MRI data from the Healthy Brain Network pediatric mental health study (***Alexander et al., 2017***), containing dMRI data from 2747 participants aged 5-21 years. These data were measured using a 1.5 T Siemens mobile scanner on Staten Island (SI) and three fixed 3 T Siemens MRI scanners at sites in the New York area: Rutgers University Brain Imaging Center (RU), the CitiGroup Cornell Brain Imaging Center (CBIC), and the City University of New York Advanced Science Research Center (CUNY). Informed consent was obtained from each participant aged 18 or older. For participants younger than 18, written consent was obtained from their legal guardians and written assent was obtained from the participant. Voxel resolution was 1.8 mm × 1.8 mm × mm with 64 non-colinear directions measured for each of *b* = 1000 s/mm^2^ and *b* = 2000 s/mm^2^.

### BIDS curation

We curated the imaging metadata for 2615 of the 2747 currently available HBN participants. Using dcm2bids and custom scripts, we conformed the data to the Brain Imaging Data Structure (BIDS; (***Gorgolewski et al., 2016***)) Specification. The BIDS-curated dataset is available on FCP-INDI and can be accessed via AWS S3 at s3://fcp-indi/data/Projects/HBN/BIDS_curated/.

After conforming the data to BIDS, we used the “Curation of BIDS” (CuBIDS) package (***Covitz et al., 2022***) to identify unique combinations, or “variants” of imaging parameters in the curated dataset. CuBIDS is a Python-based software package that provides a sanity-preserving workflow to help users reproducibly parse, validate, curate, and understand heterogeneous BIDS imaging datasets. CuBIDS includes a robust implementation of the BIDS Validator that scales to large samples and incorporates DataLad (***Halchenko et al., 2021***), a distributed data management system, to ensure reproducibility and provenance tracking throughout the curation process. CuBIDS tools also employ agglomerative clustering to identify the aforementioned variants of imaging parameters. Users may then test BIDS-Apps on a subset of participants that represent the full range of acquisition parameters that are present. These variants are listed in the participants.tsv file in the BIDS-curated dataset.

### Preprocessing

We performed dMRI preprocessing on 2615 participants, using *QSIPrep* (***Cieslak et al., 2021***) 0.12.1, which is based on *Nipype* 1.5.1 (***Gorgolewski et al., 2011***, ***2018***), RRID:SCR_002502. *QSIPrep* a robust and scalable pipeline to group, distortion correct, motion correct, denoise, coregister and resample MRI scans. In total, 417 participants failed this preprocessing step, largely due to missing dMRI files. In keeping with the BIDS Specification, the preprocessed dataset is available as a derivative dataset within the BIDS-curated dataset and can be access on AWS S3 at s3://fcp-indi/data/Projects/HBN/ BIDS_curated/derivatives/qsiprep/. *QSIPrep* fosters reproducibility by automatically generating thorough methods boilerplate for later use in scientific publications, which we use for the remainder of this subsection to document each preprocessing step.

#### • Anatomical data preprocessing

The T1-weighted (T1w) image was corrected for intensity nonuniformity (INU) using `N4BiasFieldCorrection` (***Tustison et al., 2010***) (ANTs 2.3.1), and used as T1w-reference throughout the workflow. The T1w-reference was then skull-stripped using `antsBrainExtraction.sh` (ANTs 2.3.1), using OASIS as target template. Spatial normalization to the ICBM 152 Nonlinear Asymmetrical template version 2009c (***Fonov et al. (2009b)***, RRID:SCR_008796) was performed through nonlinear registration with `antsRegistration` (***Avants et al. (2008)***, ANTs 2.3.1, RRID:SCR_004757), using brain-extracted versions of both T1w volume and template. Brain tissue segmentation of cerebrospinal fluid (CSF), white-matter (WM) and gray-matter (GM) was performed on the brain-extracted T1w using FAST (***Zhang et al., 2001***), FSL 6.0.3:b862cdd5, RRID:SCR_002823.

#### • Diffusion data preprocessing

Any images with a *b*-value less than 100 s/mm^2^ were treated as a *b* = 0 image. MP-PCA denoising as implemented in MRtrix3’s `dwidenoise` (***Veraart et al., 2016***) was applied with a 5-voxel window. After MP-PCA, B1 field inhomogeneity was corrected using `dwibiascorrect` from MRtrix3 with the N4 algorithm (***Tustison et al., 2010***). After B1 bias correction, the mean intensity of the DWI series was adjusted so all the mean intensity of the *b* = 0 images matched across each separate DWI scanning sequence.

FSL (version 6.0.3:b862cdd5)’s eddy was used for head motion correction and Eddy current correction (***Andersson and Sotiropoulos, 2016***). Eddy was configured with a *q*-space smoothing factor of 10, a total of 5 iterations, and 1000 voxels used to estimate hyperparameters. A linear first level model and a linear second level model were used to characterize Eddy current-related spatial distortion. *q*-space coordinates were forcefully assigned to shells. Field offset was attempted to be separated from participant movement. Shells were aligned post-eddy. Eddy’s outlier replacement was run (***Andersson et al., 2016***). Data were grouped by slice, only including values from slices determined to contain at least 250 intracerebral voxels. Groups deviating by more than four standard deviations from the prediction had their data replaced with imputed values. Data was collected with reversed phase-encode blips, resulting in pairs of images with distortions going in opposite directions. Here, *b* = 0 reference images with reversed phase encoding directions were used along with an equal number of *b* = 0 images extracted from the DWI scans. From these pairs the susceptibility-induced off-resonance field was estimated using a method similar to that described in (***Andersson et al., 2003***). The fieldmaps were ultimately incorporated into the Eddy current and head motion correction interpolation. Final interpolation was performed using the `jac` method. Several confounding time-series were calculated based on the *preprocessed DWI*: framewise displacement (FD) using the implementation in *Nipype* following the definitions by (***Power et al., 2014***). The DWI time-series were resampled to ACPC, and their corresponding gradient directions were rotated accordingly to generate a *preprocessed DWI run in ACPC space*.

Many internal operations of *QSIPrep* use *Nilearn* 0.6.2 (***Abraham et al., 2014***), RRID:SCR_001362 and *DIPY* (***Garyfallidis et al., 2014***). For more details of the pipeline, see the section corresponding to workflows in *QSIPrep*’s documentation.

### Cloud-based distributed preprocessing

The containerization of *QSIPrep* provided a consistent preprocessing pipeline for each participant but the number of participants made serial processing of each participant prohibitive on a single machine. We used *cloudknot*, a previously developed cloud-computing library (***Richie-Halford and Rokem, 2018***) to parallelize the preprocessing over individual participants on spot instances in the Amazon Web Services Batch service. *Cloudknot* takes as input a user-defined Python function and creates the necessary AWS infrastructure to map that function onto a range of inputs, in this case, the participant IDs. The Python preprocessing function was a thin wrapper around *QSIPrep*’s command line interface and is provided in a Jupyter notebook in the HBN-POD2 GitHub repository in the “notebooks” directory. Using *cloudknot* and AWS Batch Spot Instances, the preprocessing cost less than $1.00 per participant.

### Expert quality control

The expert QC “gold standard” subset was created by randomly selecting 200 participants from the preprocessed dataset, sampled such that the proportional site distribution in the gold standard subset matched that of the preprocessed dataset.

We created a web application for expert quality control of preprocessed dMRI, called *dmriprepviewer* (***Richie-Halford et al., 2022***). The viewer ingests *QSIPrep* outputs and generates a browser-based interface for expert QC. It provides a study overview displaying the distributions of *QSIPrep*’s automated data quality metrics (described at https://qsiprep.readthedocs.io/en/latest/preprocessing.html#quality-control-data). Each datum on the study overview page is interactively linked to a participant-level QC page that provides an interactive version of *QSIPrep*’s visual reports (described at https://qsiprep.readthedocs.io/en/latest/preprocessing.html#visual-reports). The viewer allows users to assign a rating of −2 (definitely fail), −1 (probably fail), 0 (not sure), 1 (probably pass), or 2 (definitely pass) to a participant. To standardize rater expectations before rating, expert raters watched a tutorial video (available on YouTube at https://youtu.be/SQ0v-O-e5b8 and in the OSF project). They then rated each participant and saved their scores and sent them to the lead author for compilation.

To compute the pairwise Cohen’s *κ* scores in Figure 3e, we used the *scikit-learn* (***Pedregosa et al., 2011***) `cohen_kappa_score` function with quadratic weights. To compute intra-class correlation, we used the *pingouin* statistical package (***Vallat, 2018***) `intraclass_corr` function. The expert rating analysis can be replicated using the `make expert-qc` command in the HBN-POD2 GitHub repository.

The mean expert ratings were scaled to the range 0 to 1, so that a mean rating from 0 to 0.2 corresponds to an expert rating of “definitely fail”, a mean rating from 0.2 to 0.4 corresponds to “probably fail”, from 0.4 to 0.6 corresponds to “not sure”, from 0.6 to 0.8 corresponds to “probably pass”, and 0.8 to 1.0 corresponds to “definitely pass.” These expert scores are available in the “expert_qc_score” column of the `participants.tsv` file on FCP-INDI.

### Community scientist quality control

The community science web application is based on the SwipesForScience framework https://swipesforscience.org/, which generates a web application for community science given an open repository of images to be labelled and a configuration file. The source code for the *Fibr* web application is available at https://github.com/richford/fibr. After a brief tutorial, community scientists provided binary pass/fail ratings based on the DEC-FA from a fit of a DTI model to each participant’s preprocessed dMRI data. These images were generated using a *DIPY* (***Garyfallidis et al., 2014***) `TensorModel` in a *cloudknot*-enabled Jupyter notebook that is available in the “notebooks” directory of the *Fibr* GitHub repository. *Fibr* saves each community rating to its Google Firebase backend, the contents of which have been archived to the HBN-POD2 OSF project.

The *Fibr* ratings were then combined with 31 automated *QSIPrep* data quality metrics to train the gradient boosted trees models XGB, XGB-f, and XGB-q. See Appendix 3 for a list of these automated QC metrics and a measure of their global feature importance in the XGB and XGB-q models. These models were implemented as binary classifiers using the XGBoost library (***Chen and Guestrin, 2016b***). The targets for these classifiers were the mean expert ratings in the gold standard dataset, rescaled to the range [0, 1] and binarized with a threshold of 0.5. Using repeated stratified K-fold cross-validation, with three splits and two repeats, we evaluated the models’ performance in predicting the gold standard ratings. In each fold, the best model hyperparameters were chosen using the scikit-optimize (***Head et al., 2021***) `BayesSearchCV class`. Saved model checkpoints for each crossvalidation split are available in the HBN-POD2 OSF project. Since each split resulted in a different XGB model and we required a single QC score to train the deep learning model, we combined the models from each cross-validation split using a voting classifier, computing a weighted averaged of the predicted probability of passing from each model, weighted by its out-of-sample ROC-AUC. This was implemented using scikit-learn’s `VotingClassifier class`. Treating the voting classifier as another “expert” rater, we reassessed the pairwise Cohen’s *κ* and ICC scores as in the expert QC subsection. The community ratings analysis can be replicated using the `make community-qc` command in the HBN-POD2 GitHub repository. The XGB model’s positive class probabilities are available in the “xgb_qc_score” column of the `participants.tsv` file on FCP-INDI, while the XGB-q model’s positive class probabilities are available in the “xgb_qsiprep_qc_score” column.

### Deep learning to predict quality control

The binarized voting classifier’s predictions were then used as targets to train a deep learning binary classifier to predict QC scores based on each participant’s preprocessed dMRI data. We trained two different model architectures: (i) CNN-i, which took only preprocessed dMRI images as input and (ii) CNN-i+q, whose input also included *QSIPrep*’s automated data quality metrics. Both models were implemented in Tensorflow 2 (***Abadi et al., 2015***) using the Keras module (***Chollet et al., 2015***). The image processing part of the model architecture was identical for both models: a modification of an existing 3D CNN (***Zunair et al., 2020***) previously applied to assess tuberculosis severity (***Dicente Cid et al., 2019***). It accepts a 3D volume as input with four channels: (i) the *b* = 0 reference volume, (ii) DEC-FA in the *x*-direction, (iii) DEC-FA in the *y*-direction and (iv) DEC-FA in the *z*-direction. The *QSIPrep*’s automated QC metrics were included as an additional fifth channel. The CNN-i+q model architecture is summarized in Figure 10. Upon input, the CNN-i+q model extracts the imaging channels and passes them through the CNN architecture. The remaining data quality metrics channel is flattened and passed “around” the CNN architecture and concatenated with the output of the convolutional layers. This concatenated output is then passed through a fully-connected layer to produce a single output, the probability of passing QC. This architecture has 1,438,783 trainable parameters.

**Figure 10.**
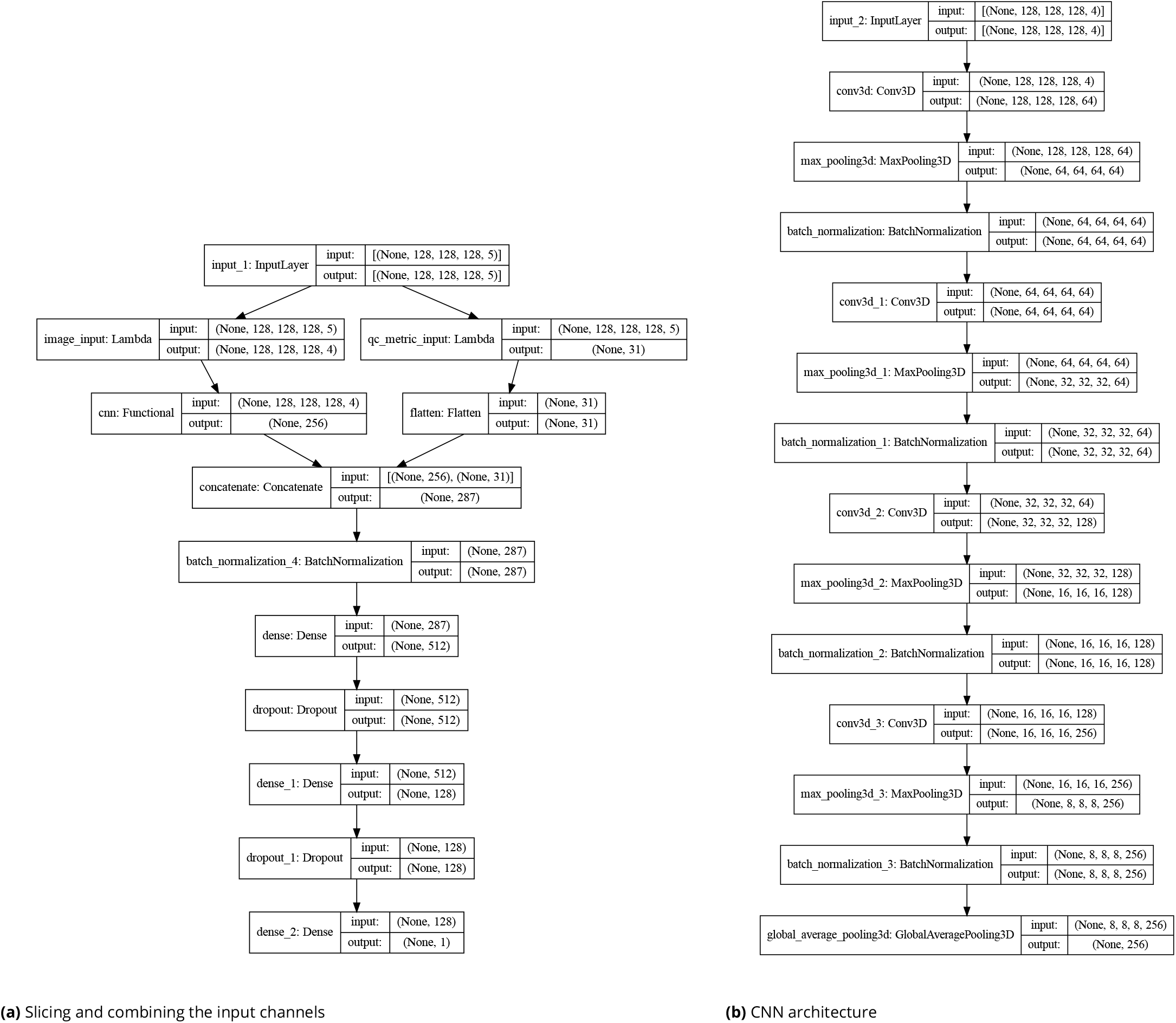
Deep learning model architecture: **(a)** The CNN-i+q model accepts multichannel input that combined four imaging channels with a fifth channel containing 31 *QSIPrep* automated data quality metrics. The imaging channels are separated from the data quality channel using Lambda layers. The imaging channels are passed through a CNN **(b)**, the output of which is concatenated with the data quality metrics, batch normalized and passed through two fully-connected (FC) layers, with rectified linear unit (ReLu) activation functions and with 512 and 128 units respectively. Each FC layer is followed by a dropout layer which drops 40% of the input units. The final layer contains a single unit with a sigmoid activation function and outputs the probability of passing QC. **(b)** The CNN portion of the model passes the imaging input through four convolutional blocks. Each block consists of a 3D convolutional layer with a kernel size of 3 and a ReLu activation, a 3D max pooling layer with a pool size of 2, and a batch normalization layer with Tensorflow’s default parameters. The number of filters in the convolutional layers in each block are 64, 64, 128, and 256 respectively. The output of the final block is passed through a 3D global average pooling layer with Tensorflow’s default parameters. **Figure 10–Figure supplement 1.** Deep learning model loss curves

We used *DIPY* (***Garyfallidis et al., 2014***) and *cloudknot* (***Richie-Halford and Rokem, 2018***) to generate these multichannel volumes for each participant and save them as NIfTI-1 files (***Cox et al., 2004***). These NIfTI files were then converted to the Tensorflow TFRecord format using the *Nobrainer* deep learning framework (***Kaczmarzyk et al., 2021***). The Jupyter notebooks used to create these NIfTI and TFRecord files are available in the “notebooks” directory of the *Fibr* GitHub repository.

We trained each model using the Google Cloud AI Platform Training service; the HBN-POD2 GitHub repository contains Docker services to launch training (with `make dl-train`) and prediction (with `make dl-predict`) jobs on Google Cloud, if the user has provided the appropriate credentials in an environment file and placed the TFRecord files on Google Cloud Storage. To estimate the variability in model training, we trained ten separate models using different training and validation splits of the data. The gold standard dataset was not included in any of these splits and was reserved for reporting final model performance. Models were optimized for binary crossentropy loss using the Adam optimizer (***Kingma and Ba, 2017***) with an initial learning rate of 0.0001. We reduced the learning rate by a factor of 0.5 when the validation loss plateaued for more than two epochs. We also stopped training when the validation loss failed to improve by more than 0.001 for twenty consecutive epochs. These two adjustments were made using the `ReduceLROnPlateau` and `EarlyStopping` callbacks in Tensorflow 2 (***Abadi et al., 2015***) respectively. The training and validation loss curves for both the CNN-i and CNN-i+q models are depicted in ***Figure 10–Figure Supplement 1***. While the CNN-i+q model achieved better validation loss, it did not outperform the CNN-i model on the held out gold standard dataset. The CNN-i+q model’s positive class probabilities are available in the “dl_qc_score” column of the `participants.tsv` file on FCP-INDI.

To generate the attribution maps, we followed Tensorflow’s integrated gradients tutorial (***TensorFlow Authors, 2021***) with a black baseline image and 128 steps in the Riemann sum approximation of the integral (i.e. `m_steps = 128`). In the HBN-POD2 GitHub repository, we provide a Docker service to compute integrated gradient attribution maps on Google Cloud, which can be invoked using the make `dl-integrated-gradients` command.

### Site generalization experiments

To simulate the generalization of the XGB-q and CNN-i models to new scanning sites, we trained multiple versions of XGB-q and CNN-i with different scanning sites held out and then evaluated those models on the held out sites. These models were therefore evaluated on data from “unseen” sites. We constructed these train/evaluate splits from combinations of the HBN sites with 3 T scanners (RU, CBIC, and CUNY), and excluded CUNY as a standalone training or test site because of its low number of participants (*N* = 74). This left four combinations of site-generated training splits: CBIC + CUNY (eval: RU), CBIC (eval: RU + CUNY), RU + CUNY (eval: CBIC), and RU (eval: CBIC + CUNY).

We trained eight models (with distinct random seeds) from the CNN-i family of models using the global XGB scores as targets, just as with the full CNN-i model. Similarly, we trained twenty models (with distinct random seeds) from the XGB-q family of models using the expert scores as targets, just as with the full XGB-q model. For each model, we reported three evaluation metrics: ROC-AUC, accuracy, and balanced accuracy. Because the distribution of QC scores was imbalanced (Figures 3a and 5d), we included balanced accuracy as an evaluation metric. Balanced accuracy avoids inflated accuracy estimates on imbalanced data (***Velez et al., 2007***), and in the binary classification case, it is the mean of the sensitivity and specificity. For the CNN-i family, we further decomposed the evaluation split into a report set, for which expert scores were available, and a test set, with participants who were not in the “gold standard” dataset. For the report set, we evaluated the model using the expert scores as the ground truth. For the test set, we evaluated each model using the XGB scores as ground truth.

Aside from the Specification of train and evaluation splits, model training followed exactly the same procedure as for the full dataset. For example, we use the same cross validation and hyperparameter optimization procedure for the XGB-q family as for the original XGB-q model and the same architecture, input format, and early stopping criteria for the CNN-i family as for the CNN-i model. In the HBN-POD2 GitHub repository, we provide a Docker service to conduct the CNN-i site generalization experiments Google Cloud, which can be invoked using the `make dl-site-generalization` command. The XGB-q site generalization experiments can be replicated locally using the make `site-generalization` command, which will also plot the results of the CNN-i experiments.

### QC bundle profiles

To generate bundle profiles, reconstruction was performed using the *QSIprep* 0.12.1 preconfigured reconstruction workflow `mrtrix_multishell_msmt`, modified to generate two million streamlines rather than the default ten million. Multi-tissue fiber response functions were estimated using the dhollander algorithm. Fiber orientation distributions (FODs) were estimated via constrained spherical deconvolution (CSD, (***Tournier et al., 2004***, ***2008***)) using an unsupervised multi-tissue method (***Dhollander et al., 2019***, ***2016***). Reconstruction was done using MRtrix3 (***J-Donald et al., 2019***). FODs were intensity-normalized using mtnormalize (***Raffelt et al., 2017***).

These tractograms were then used as input to the Python Automated Fiber Quantification toolbox (pyAFQ) (***Kruper et al., 2021***). Twenty-four major tracts, which are enumerated in Figure 8, were identified using multiple criteria: streamlines are selected as candidates for inclusion in a bundle of streamlines that represents a tract if they pass through known inclusion ROIs and do not pass through exclusion ROIs (***Wakana et al., 2007***). In addition, a probabilistic atlas is used to exclude streamlines which are unlikely to be part of a tract and to adjudicate in cases where a streamline could belong to more than one tract (***Hua et al., 2008***). Each streamline is resampled to 100 nodes and the robust mean at each location is calculated by estimating the 3D covariance of the location of each node and excluding streamlines that are more than 5 standard deviations from the mean location in any node. Finally, a bundle profile of tissue properties in each bundle was created by interpolating the value of MRI maps of these tissue properties to the location of the nodes of the resampled streamlines designated to each bundle. In each of 100 nodes, the values are summed across streamlines, weighting the contribution of each streamline by the inverse of the mahalanobis distance of the node from the average of that node across streamlines. This means that streamlines that are more representative of the tract contribute more to the bundle profile, relative to streamlines that are on the edge of the tract.

These processes create bundle profiles, in which diffusion measures are quantified and averaged along twenty-four major fiber tracts. We retain only the mean diffusivity (MD) and the fractional anisotropy (FA) from a diffusion kurtosis imaging (DKI) model (***Jensen et al., 2005***), implemented in DIPY (***Henriques et al., 2021***), and impute missing bundles using median imputation as implemented by *scikit-learn*’s `SimpleImputer class`. Because the HBN-POD2 bundle profiles exhibit strong site effects (***Richie-Halford et al., 2021***), we used the ComBat harmonization method to robustly adjust for site effects in the tract profiles. Initially designed to correct for site effects in gene expression studies (***Johnson et al., 2007***), ComBat employs a parametric empirical Bayes approach to adjust for batch effects and has since been applied to multi-site cortical thickness measurements (***Fortin et al., 2018***), multi-site DTI studies (***Fortin et al., 2017***), and functional MRI data in the Adolescent Brain Cognitive Development Study (ABCD) (***Nielson et al., 2018***). We rely on the *neurocombat_sklearn* library (***Pinaya, 2020***), to apply ComBat in before plotting bundle profiles in Figure 8 using plotting functions from the AFQ-Insight package (***Richie-Halford et al., 2019***). The bundle profile analysis can be replicated using the `make bundle-profiles` command in the HBN-POD2 GitHub repository.

### Brain age prediction

We evaluated the effect of varying the QC cutoff on model performance by observing cross-validated *R*^2^ values of gradient boosted trees models implemented using XGBoost. The input feature space for each model consisted of 4800 features per participant, comprising 100 nodes for each of MD and FA in the twenty-four major tracts. We imputed missing bundles and harmonized the different scanning sites as above. The XGBoost models’ hyperparameters were hand-tuned to values that have been performant in the authors’ previous experience. Within the limited age range of the HBN study, MD and FA follow logarithmic maturation trajectories (***Yeatman et al., 2014***). We therefore log-transformed each participant’s age before prediction using the `TransformedTargetRegressor` class from *scikit-learn*. For each value of the QC cutoff between 0 and 0.95, in steps of 0.05, we computed the cross-validated *R*^2^ values using *scikit-learn*’s `cross_val_score` function with repeated K-fold cross-validation using five folds and five repeats.

## Author contributions statement

The last two authors named share senior authorship. The first two authors named share lead authorship. The remaining authors are listed in alphabetical order, with the exception of the *Fibr* Community Science Consortium, whose members provided community science QC ratings and are listed in Appendix 4. We describe contributions to the paper using the CRediT taxonomy (***Brand et al., 2015***; ***Allen et al., 2014***): Conceptualization: A.R-H., A.R., T.S., and M.C.; Methodology: A.R-H. and A.R.; Software: A.R-H., M.C., and S.C.; Validation: A.R-H., M.C., and S.C.; Formal Analysis: A.R-H. and M.C.; Investigation: A.R-H. and M.C.; Resources: A.R., T.S., and M.M.; Data Curation: S.C., M.C., V.J.S., I.I.K., B.A-P. and L.A.; Writing – Original Draft: A.R-H. and A.R.; Writing – Review & Editing: A.R-H., A.R., M.C., A.F., T.S., V.J.S., I.I.K, B.A-P., and S.C.; Visualization: A.R-H.; Supervision: A.R. and T.S.; Project Administration: A.R-H. and A.R.; Funding Acquisition: A.R. and T.S.

## Acknowledgments

We would like to thank Anisha Keshavan for useful discussions of community science and web-based quality control and for her work on SwipesForScience. This manuscript was prepared using a limited access dataset obtained from the Child Mind Institute Biobank, The Healthy Brain Network dataset. This manuscript reflects the views of the authors and does not necessarily reflect the opinions or views of the Child Mind Institute. This work was supported via BRAIN Initiative grant 1RF1MH121868-01 from the National Institutes of Mental Health. Additional support was provided by grant 1R01EB027585-01 from the National Institutes of Biomedical Imaging and Bioengineering (PI: Eleftherios Garyfallidis). Additional support was provided by R01MH120482 and the Penn/CHOP Lifespan Brain Institute.

## Appendix 1 CuBIDS variant annotation

We identified 20 unique dMRI acquisitions across HBN-POD2, which are summarized in Table 1. Site CBIC has two acquisition types: “64dir,” which shares it’s pulse sequence with sites RU and CUNY, and “ABCD64dir,” with acquisition parameters that better match the ABCD study (TE=0.089 s and TR=4.1 s). The “Most_Common” variant identifies the most common combination of acquisition parameters for a given site and acquisition. The “Low_Volume” variant identifies participants from all sites with less that 129 DWI volumes, which is the number of volumes in the most common variants. All remaining variants names identify the acquisition parameter(s) that differ from those of the most common variant. For example, the “MultibandAccelerationFactor” variant has a different multiband acceleration factor than that of the the most common variant but all participants within that variant share the same multiband acceleration factor. Variants that differ by multiple acquisition parameters have names that are composed of concatenated parameters. For example, the variant “Dim3SizeVoxelSizeDim3” varies both in the number of voxels in dimension 3 (“Dim3Size”) and in the voxel size in dimension 3 (“VoxelSizeDim3”).

**Appendix 1 Table 1.** Participant counts for HBN-POD2 variants.

| Site | Acquisition | Variant | Count |
| --- | --- | --- | --- |
| CBIC | 64dir | Most_Common | 828 |
| CBIC | 64dir | Obliquity | 32 |
| CBIC | 64dir | VoxelSizeDim1VoxelSizeDim2 | 1 |
| CBIC | ABCD64dir | Most_Common | 15 |
| CBIC | ABCD64dir | HasFmap | 2 |
| CBIC | ABCD64dir | MultibandAccelerationFactor | 1 |
| CBIC | ABCD64dir | Obliquity | 1 |
| CUNY | 64dir | Most_Common | 68 |
| CUNY | 64dir | Dim3SizeVoxelSizeDim3 | 4 |
| CUNY | 64dir | Obliquity | 2 |
| RU | 64dir | Most_Common | 859 |
| RU | 64dir | NoFmap | 5 |
| RU | 64dir | Obliquity | 8 |
| RU | 64dir | PhaseEncodingDirection | 1 |
| SI | 64dir | EchoTime | 1 |
| SI | 64dir | EchoTimePhaseEncodingDirection | 9 |
| SI | 64dir | Most_Common | 269 |
| SI | 64dir | NoFmap | 2 |
| SI | 64dir | Obliquity | 12 |
| All Sites | All Acquisitions | Low_Volume_Count | 14 |

## Appendix 2 HBN-POD2 quality control instruments

We created quality control web applications for both community raters and expert raters. These apps are publicly accessible at https://fibr.dev, for the community science instrument and at http://www.nipreps.org/dmriprep-viewer/ for the expert rating instrument. We encourage readers to try these web applications on their own but have included screenshots and a summary of the interfaces in Figure 1.

**Appendix 2 Figure 1.**
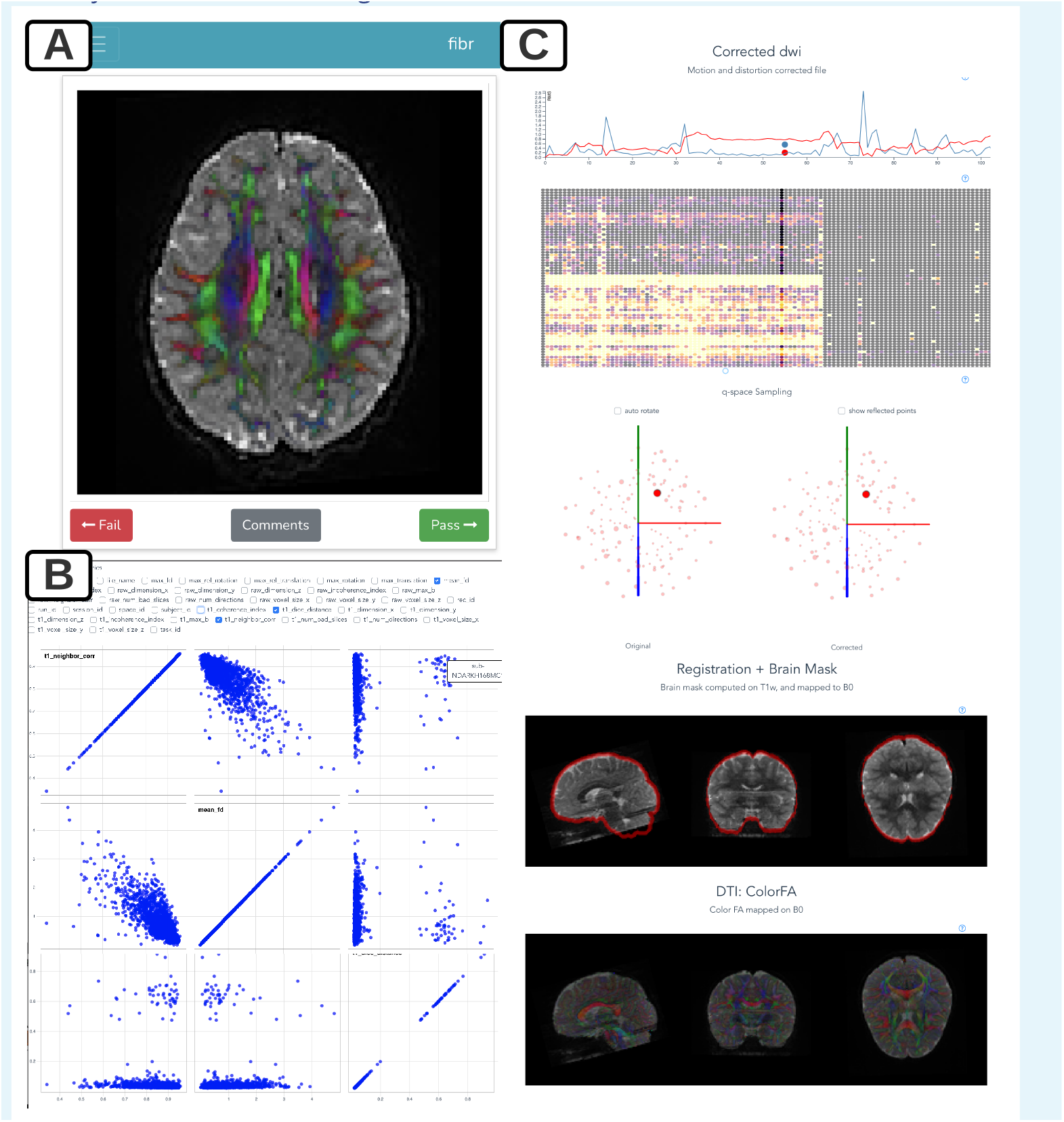
HBN-POD2 quality control instruments: **(A)** The user interface for community science QC app *Fibr*. After a tutorial, users are asked to give binary pass/fail ratings to each subject’s DEC-FA image. The intuitive swipe or click interface allows community scientists to review more images than is practical for expert reviewers. Expert reviewers use the more advanced *dmriprep-viewer* interface, where they can **(B)** view the distribution of data quality metrics for the entire study using interactive scatterplots and violin plots, and **(C)** inspect individual participants’ preprocessing results, including corrected dMRI images, frame displacement, q-space sampling distributions, registration information, and a DTI model.

## Appendix 3 XGB feature importance

SHAP is a method to explain individual predictions based on game theoretically optimal Shapley values (***Lundberg and Lee, 2017***). To estimate global feature importance for the XGB and XGB-q models, we use the `shap` library’s `TreeExplainer` (***Lundberg et al., 2020***) and average the absolute Shapley value per feature across each individual prediction. Tables 1 and 2 list the *QSIPrep* automated QC metric features in order of decreasing mean absolute shap value for the XGB and XGB-q models, respectively. We chose the top three metrics from Table 1 to plot metric distributions in Figure 2 and correlations with the expert QC results in Figure 3.

**Appendix 3 Table 1.**
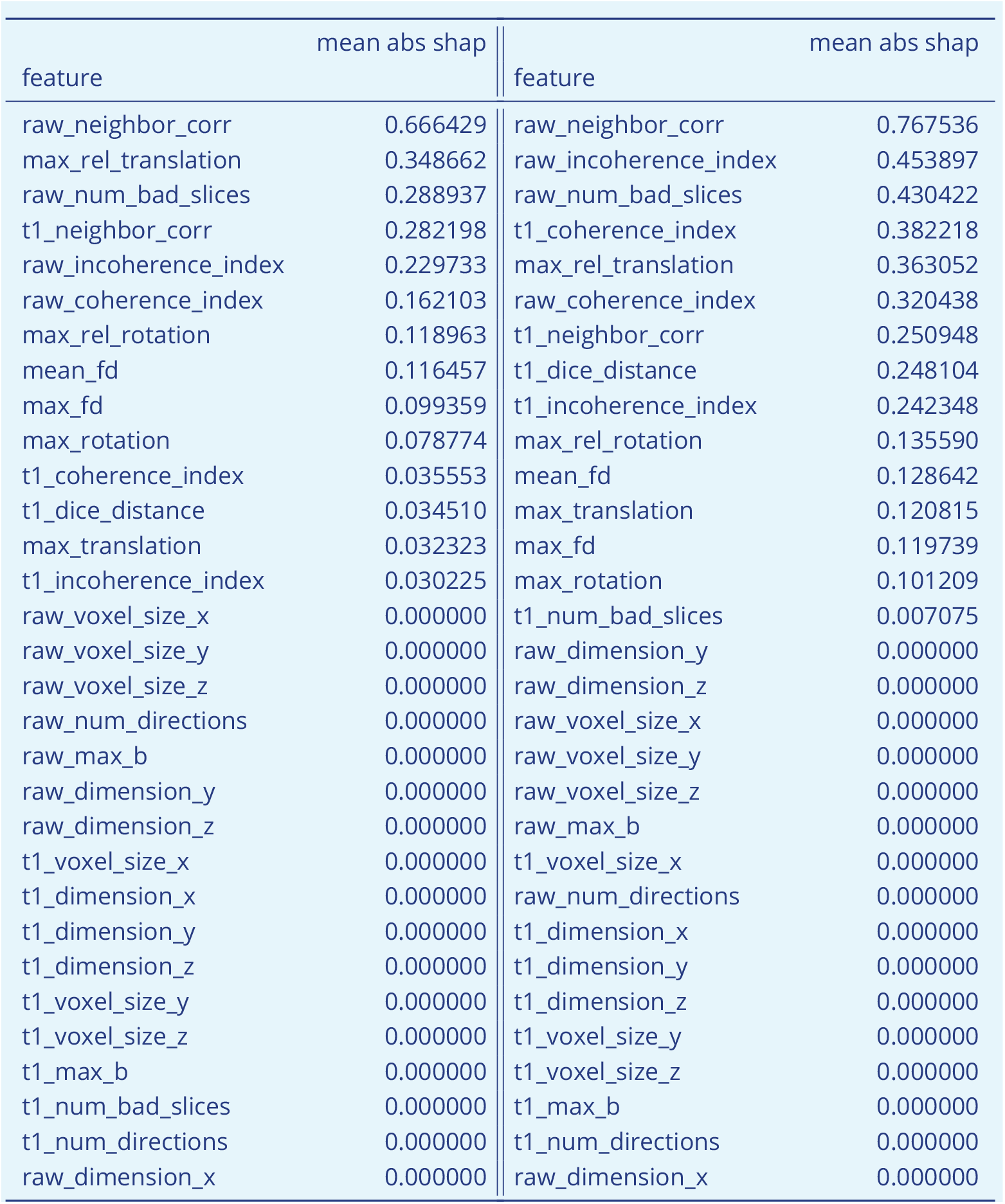
Participant counts for HBN-POD2 variants. Participant counts for HBN-POD2 variants.

| feature | mean abs shap | feature | mean abs shap |
| --- | --- | --- | --- |
| raw_neighbor_corr | 0.666429 | raw_neighbor_corr | 0.767536 |
| max_rel_translation | 0.348662 | raw_incoherence_index | 0.453897 |
| raw_num_bad_slices | 0.288937 | raw_num_bad_slices | 0.430422 |
| t1_neighbor_corr | 0.282198 | t1_coherence_index | 0.382218 |
| raw_incoherence_index | 0.229733 | max_rel_translation | 0.363052 |
| raw_coherence_index | 0.162103 | raw_coherence_index | 0.320438 |
| max_rel_rotation | 0.118963 | t1_neighbor_corr | 0.250948 |
| mean_fd | 0.116457 | t1_dice_distance | 0.248104 |
| max_fd | 0.099359 | t1_incoherence_index | 0.242348 |
| max_rotation | 0.078774 | max_rel_rotation | 0.135590 |
| t1_coherence_index | 0.035553 | mean_fd | 0.128642 |
| t1_dice_distance | 0.034510 | max_translation | 0.120815 |
| max_translation | 0.032323 | max_fd | 0.119739 |
| t1_incoherence_index | 0.030225 | max_rotation | 0.101209 |
| raw_voxel_size_x | 0.000000 | t1_num_bad_slices | 0.007075 |
| raw_voxel_size_y | 0.000000 | raw_dimension_y | 0.000000 |
| raw_voxel_size_z | 0.000000 | raw_dimension_z | 0.000000 |
| raw_num_directions | 0.000000 | raw_voxel_size_x | 0.000000 |
| raw_max_b | 0.000000 | raw_voxel_size_y | 0.000000 |
| raw_dimension_y | 0.000000 | raw_voxel_size_z | 0.000000 |
| raw_dimension_z | 0.000000 | raw_max_b | 0.000000 |
| t1_voxel_size_x | 0.000000 | t1_voxel_size_x | 0.000000 |
| t1_dimension_x | 0.000000 | raw_num_directions | 0.000000 |
| t1_dimension_y | 0.000000 | t1_dimension_x | 0.000000 |
| t1_dimension_z | 0.000000 | t1_dimension_y | 0.000000 |
| t1_voxel_size_y | 0.000000 | t1_dimension_z | 0.000000 |
| t1_voxel_size_z | 0.000000 | t1_voxel_size_y | 0.000000 |
| t1_max_b | 0.000000 | t1_voxel_size_z | 0.000000 |
| t1_num_bad_slices | 0.000000 | t1_max_b | 0.000000 |
| t1_num_directions | 0.000000 | t1_num_directions | 0.000000 |
| raw_dimension_x | 0.000000 | raw_dimension_x | 0.000000 |

## Appendix 4 The *Fibr* Community Science Consortium

The following community raters provided > 3; 000 ratings each and elected to be included in the *Fibr* Community Science Consortium as co-authors on this paper.

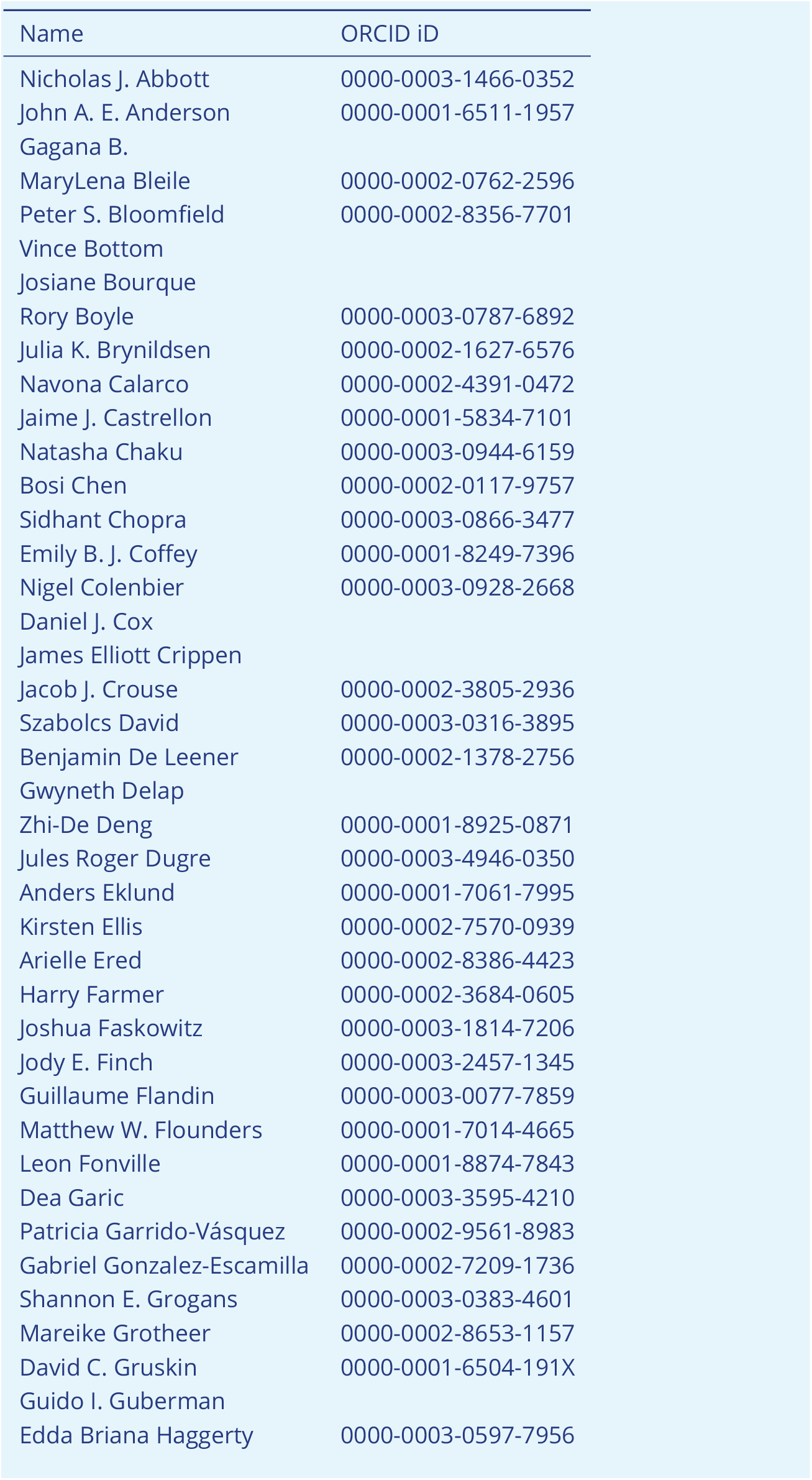
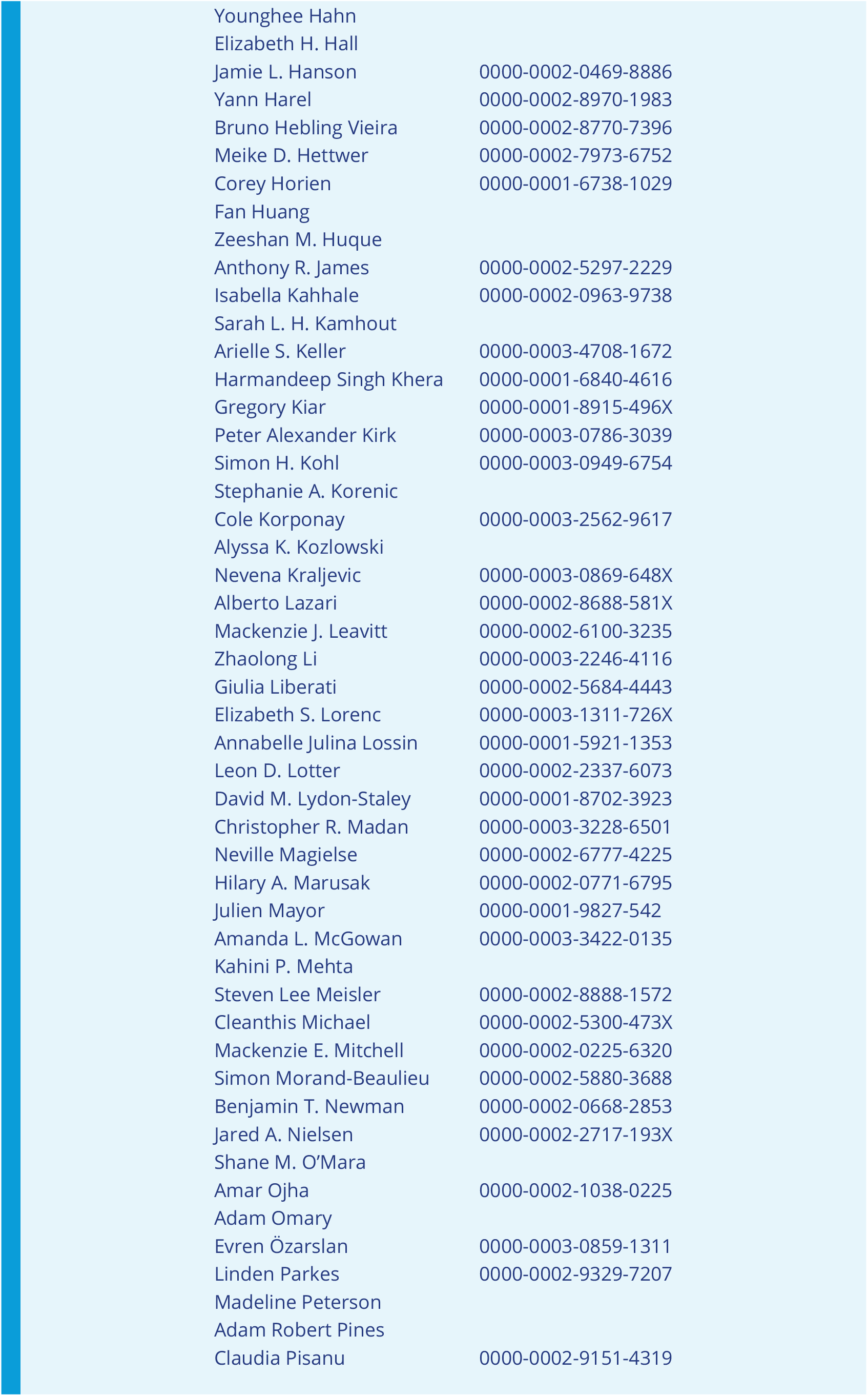
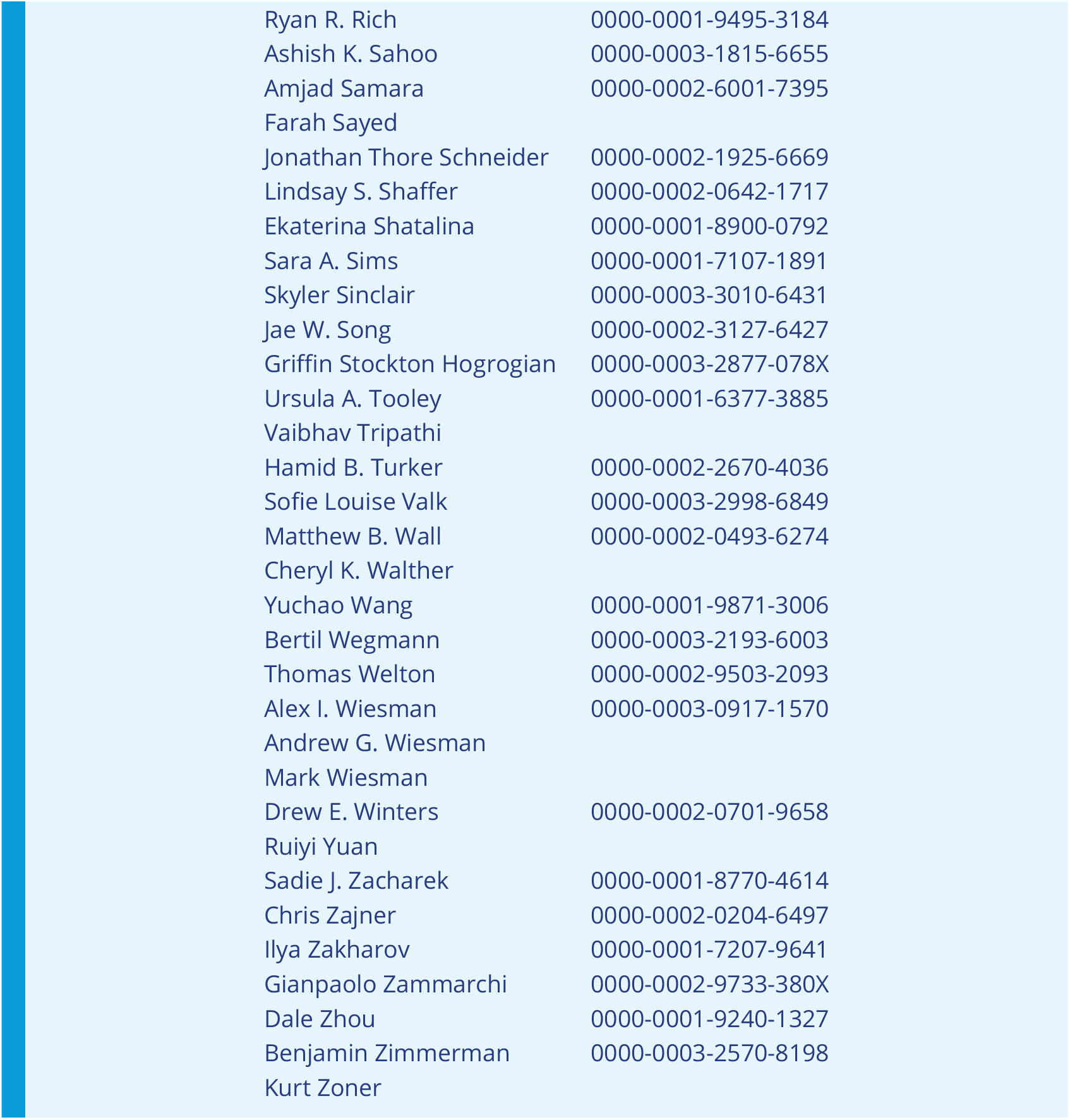

**Figure 8–Figure supplement 1.**
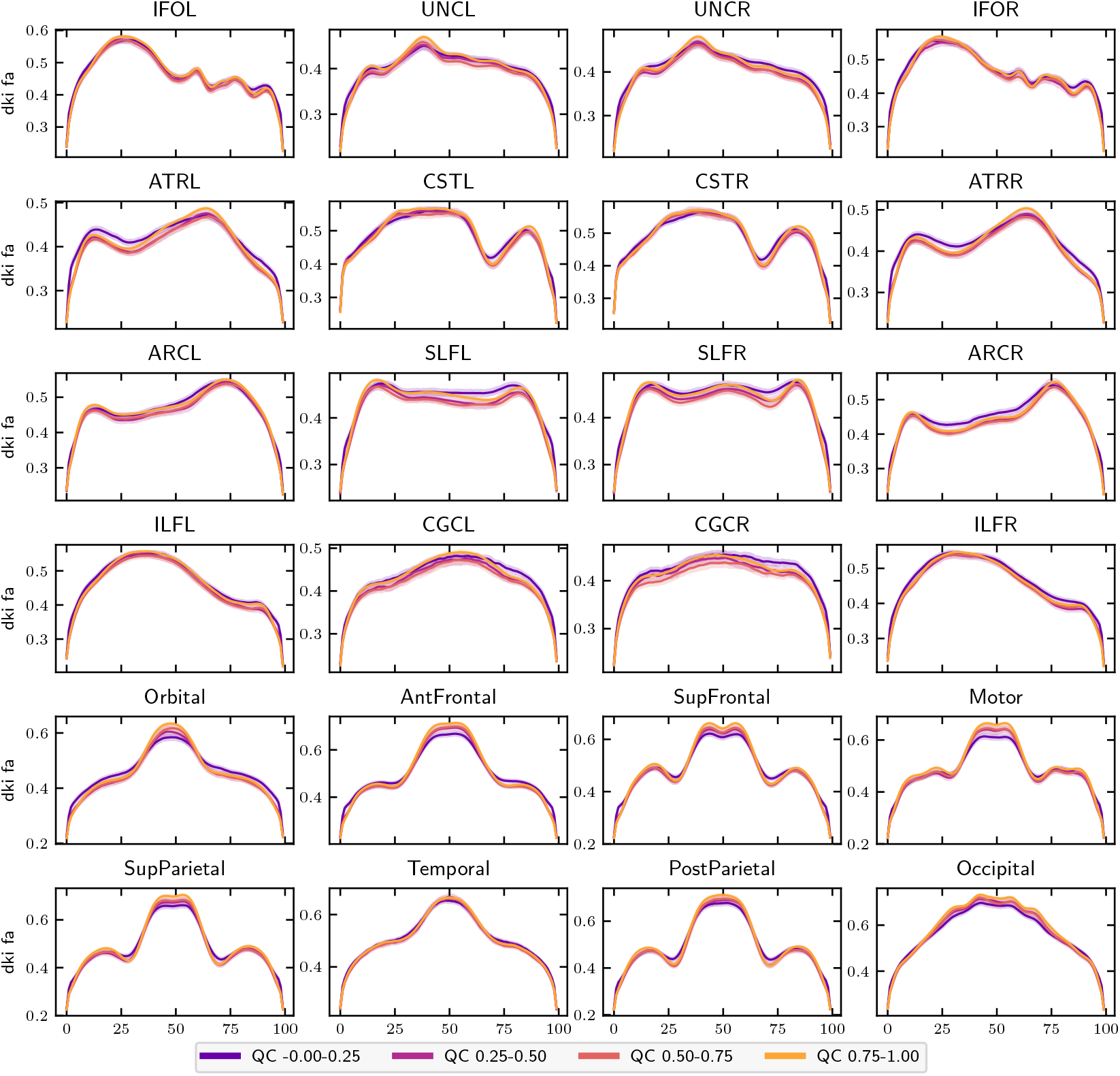
FA bundle profiles binned by QC score: FA profiles binned by QC score in twenty-four major while matter bundles. The *x*-axis represents distance along the length of the fiber bundle. Error bands represent bootstrapped 95% confidence intervals. Bundle abbreviations are as in Figure 8

**Figure 10–Figure supplement 1.**
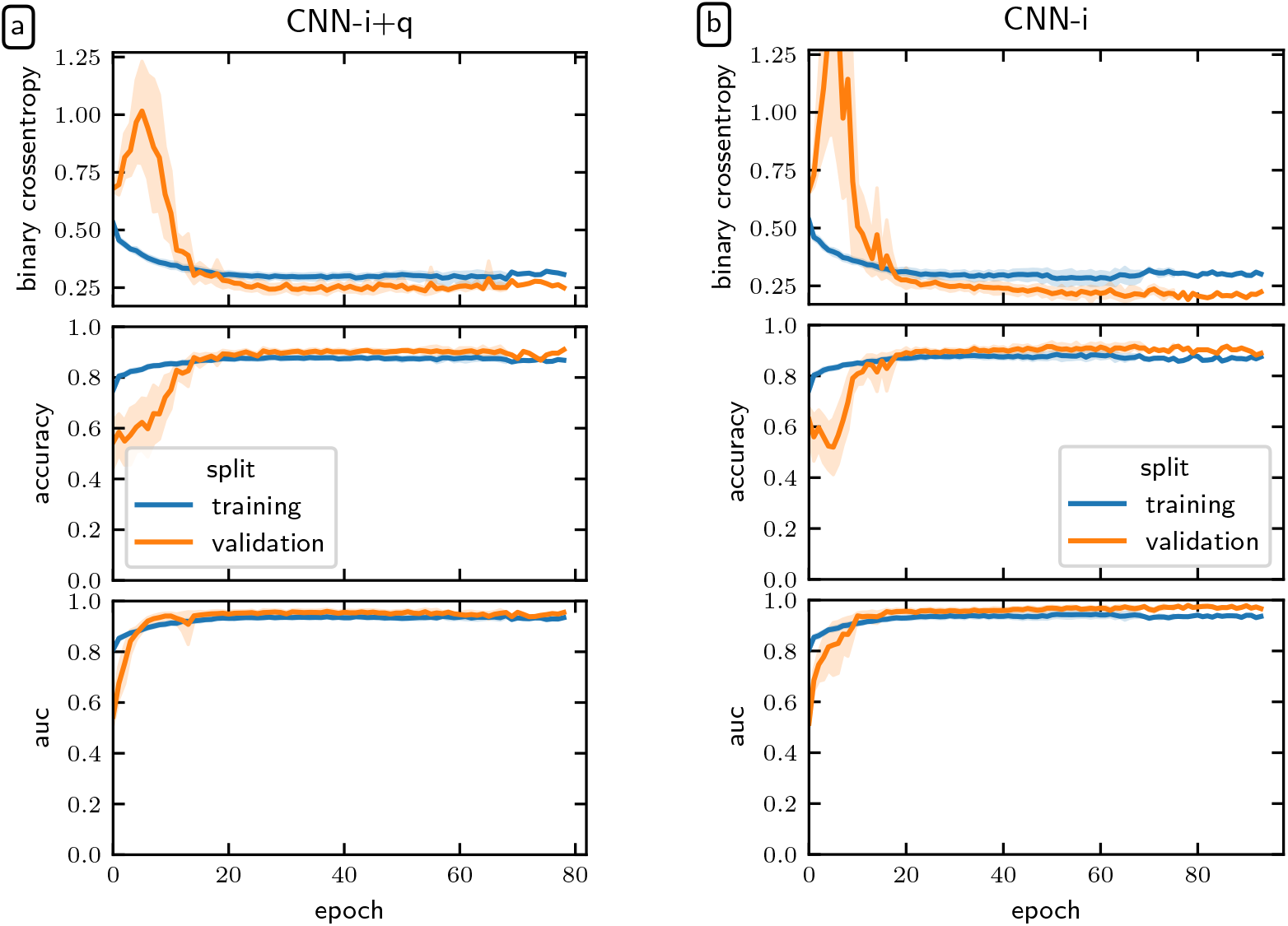
Deep learning model loss curves: The binary cross-entropy loss (top), accuracy (middle), and ROC-AUC (bottom) for (a) the CNN-i+q model and (a) the CNN-i model. Model performance typically plateaued after twenty epochs but was allowed continue until meeting the early stopping criterion. The error bands represent a bootstrapped 95% confidence interval.

## Footnotes

1 While the term “citizen science” evokes a sense of civic duty in scientific engagement, it can also imply a barrier for community members who want to contribute to science but may not be citizens of a particular country. In this manuscript we use the more modern term “community science.”

2 Curated BIDS data is available at s3://fcp-indi/data/Projects/HBN/BIDS_curated/ and *QSIPrep* outputs are available at s3://fcp-indi/data/Projects/HBN/BIDS_curated/derivatives/qsiprep/.

3 Responses for the sex variable in HBN phenotypic data are limited to “male” and “female.”

